# The DND1–NANOS3 complex shapes the primordial germ cell transcriptome via a heptanucleotide sequence in mRNA 3′ UTRs

**DOI:** 10.1101/2025.09.25.678639

**Authors:** Masataka Suzawa, Chen Qiu, Ahsan H. Polash, Misaki Yamaji, Alexis A. Jacob, Wataru Horikawa, Eva M. Farroha, Murchana Barua, Xuesong Feng, Chengyu Liu, Jason G. Williams, Davide Randazzo, Vittorio Sartorelli, Eugene Valkov, Masashi Yamaji, Traci M Tanaka Hall, Markus Hafner

## Abstract

The RNA-binding proteins DND1 and NANOS3 are essential for primordial germ cell survival^1-5^. Their co-immunoprecipitation and overlapping loss-of-function phenotypes suggest joint function^6-8^, yet how they co-regulate target mRNAs remains unclear. Here, we developed Tandem PAR-CLIP and identified a DND1–NANOS3 ribonucleoprotein that specifically recognizes an AUGAAUU heptanucleotide on target mRNAs, termed the NANOS3-dependent DND1 Recognition Element (N3-DRE). mRNAs containing 3′-UTR N3-DREs are aberrantly upregulated in DND1- or NANOS3-deficient germ cells and encode key cell-cycle and epigenome regulators, such as CDK1. Genome editing showed that the N3-DRE is essential for Cdk1 repression in mouse PGCs *in vivo*. A 1.7-Å crystal structure of the ternary complex of DND1, NANOS3, and CDK1- N3-DRE RNA revealed a continuous RNA-binding surface that confers high-affinity, sequence-specific recognition. Together, these findings define the molecular and functional basis of N3-DRE-mediated mRNA regulation in germ cell development. Moreover, we provide a paradigm of two RNA-binding proteins with low (DND1) or no (NANOS3) intrinsic sequence-specificity, jointly building a high-information-content RNA sequence motif that is different from the sum of their individual preferences. Because RNA-binding protein specificities are typically studied individually^9-13^, rather than in the context of ribonucleoproteins, this type of “two-factor authorization” may be an underappreciated mechanism to protect posttranscriptional gene regulatory networks from aberrant expression of an individual ribonucleoprotein component.

## Main

Germ cells ensure reproduction and heredity. In mammals, they originate as primordial germ cells (PGCs), the embryonic precursors of sperm and oocytes^14-18^. The fate of the PGC is specified during gastrulation from the pluripotent epiblast and then migrates toward the developing gonads, ultimately differentiating into spermatogonial stem cells (SSCs) in the testes and oogonia in the ovaries, which leads to gametogenesis in adults^14-22^. During migration, PGCs undergo developmental and genome-wide epigenetic reprogramming, characterized by cell cycle arrest at the G2/M phase, silencing of mRNA transcription, and a global shift from histone H3 lysine 9 dimethylation (H3K9me2) to lysine 27 trimethylation (H3K27me3) to confer eventual totipotency^17,20,23^.

Post-transcriptional gene regulatory (PTGR) mechanisms play an essential role in germ cell development, and germ cells exhibit a unique enrichment of cell-type-specific RNA-binding proteins (RBPs), which are often evolutionarily conserved and bind to mRNAs in a sequence-specific manner^24-26^. Disruption of these germline-specific RBPs often results in germ-cell loss and infertility^26^. One such RBP is the vertebrate-conserved DND1, which is expressed in PGCs just after specification (E6.5-6.75 in mouse) and is essential for PGC survival during their migration to the gonads^3,5,27-29^. We previously demonstrated that DND1 is essential for spermatogenesis, promoting self-renewal and preventing apoptosis of murine SSCs by recruiting the CCR4-NOT deadenylase complex to suppress key mRNAs involved in these processes^4^. DND1 is associated with another deeply conserved RBP family that also recruits CCR4-NOT, the NANOS proteins^6-8,30,31^. Mammals possess three paralogues, NANOS1-3, which differ in expression patterns and function^32,33^. NANOS3 is induced in PGCs around the same time (∼E7.5) as DND1, and its loss also results in germ-cell depletion^2,14^. The interaction of NANOS3 with DND1 and the overlapping phenotypes of *Dnd1* or *Nanos3* knockout (KO) in mice^2,5,34^ suggest a joint mechanism of gene regulation. Indeed, in human PGC-like cells (PGCLCs) differentiated from embryonic stem cells (ESCs), DND1 co-expression alters NANOS3 target mRNA selection^8^. We hypothesize that formation of a DND1–NANOS3 ribonucleoprotein (RNP) confers distinct target specificity that is essential for DND1-mediated gene regulation in germ cell development.

### Tandem fPAR-CLIP identifies a DND1–NANOS3-specific 3′ UTR motif

To comprehensively identify the RNA-binding sites of DND1, NANOS3, or the DND1–NANOS3 complex at nucleotide resolution, we used fluorescence-based PAR-CLIP (fPAR-CLIP)^35,36^. We and others previously showed that DND1-mediated target gene repression is recapitulated by ectopically expressing DND1 in HEK293 cells^4,7^, a somatic cell line that does not express germline-specific RBPs, such as DND1 or NANOS3. First, we established HEK293 cell lines expressing doxycycline (dox)-inducible N-terminally FLAG- and hemagglutinin (HA)-tagged DND1 (FH-DND1), C-terminally V5- and HA-tagged NANOS3 (NANOS3-VH), or both (Extended Data Fig. 1a). To approximate stoichiometric co-expression, FH-DND1 and NANOS3- VH were translated as a fusion protein linked by a P2A self-cleaving peptide. Dox induction resulted in robust and comparable expression levels across all lines (Extended Data Fig. 1b), and as expected, FH-DND1 and NANOS3-VH co-localized in cytoplasmic foci (Extended Data Fig. 1c).

In our co-expression cell line, we observed robust co-immunoprecipitation (co-IP) of DND1 and NANOS3. Nevertheless, considerable amounts of free DND1 and NANOS3 were also present (Extended Data Fig. 1d, e). Therefore, to specifically isolate RNA segments bound by and crosslinked to the DND1–NANOS3 RNP, we developed Tandem fPAR-CLIP by introducing an optimized tandem IP step (Extended Data Fig. 1f) that comprised sequential NANOS3-VH pulldown and FH-DND1 capture (Extended Data Fig. 1g-j). We performed three biological replicates each of fPAR-CLIP for DND1 and NANOS3 and five biological replicates of Tandem fPAR-CLIP for the DND1–NANOS3 complex (Supplementary Table 1). Spearman correlation analysis of binding sites from the biological replicates highlighted excellent reproducibility (Fig. 1a). DND1–NANOS3 Tandem fPAR-CLIP experiments formed distinct clusters on the correlation heatmap (Fig. 1a), suggesting that complex formation changed the binding specificities of DND1 and NANOS3, as suggested recently^8^.

**Figure 1.**
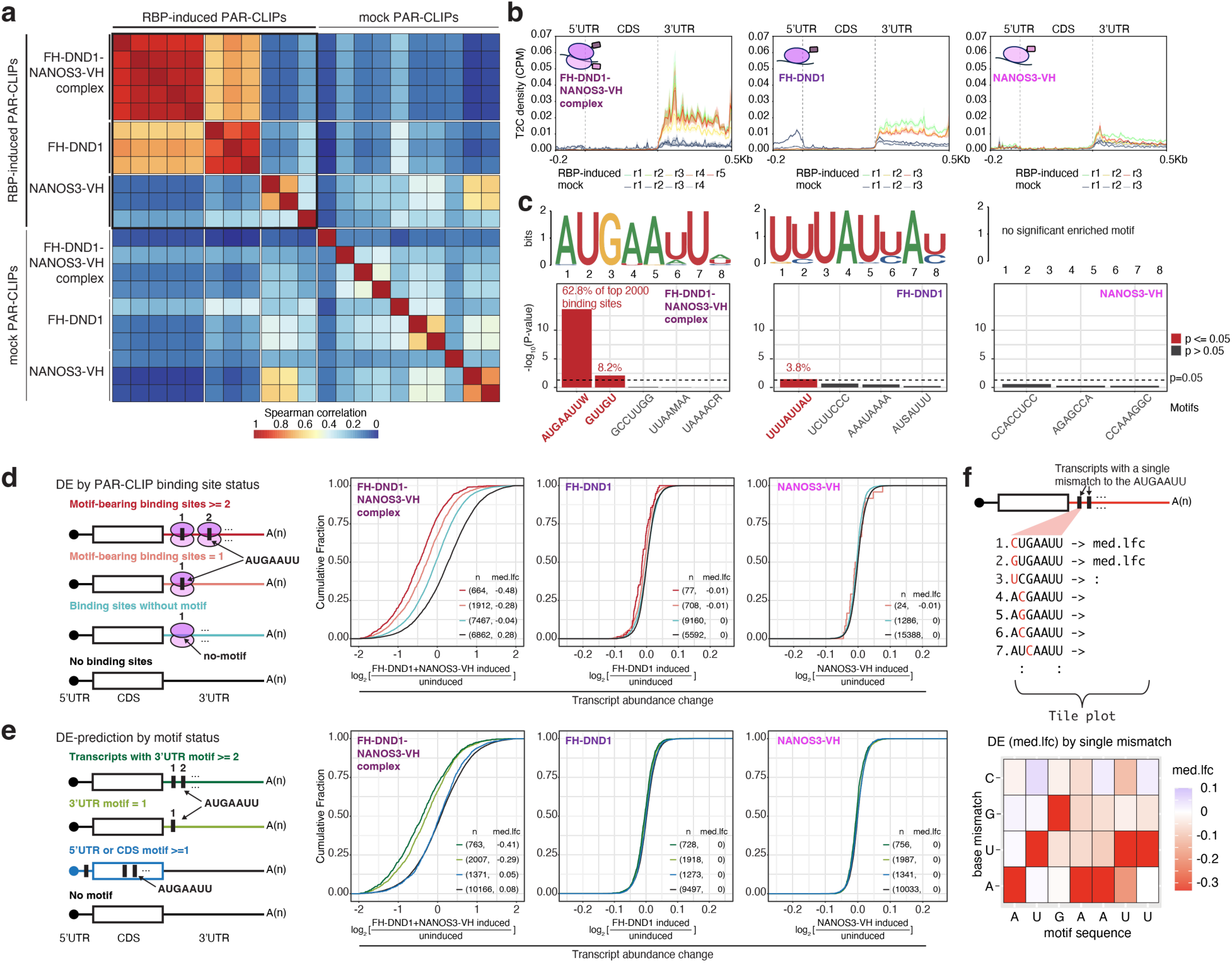
The DND1-NANOS3 complex binds and suppresses the genes with a specific sequence motif in their 3′ UTR. **(a)** Spearman correlation heatmap of all PAR-CLIP experiments based on crosslinked reads per million (XRPM) per gene. **(b)** Metagene plot of crosslinked read distribution from Tandem PAR-CLIP of the DND1–NANOS3 complex (left), or PAR-CLIP of DND1 (middle), or NANOS3 (right). **(c)** Binding motif from the top 2000 binding sites of DND1, NANOS3, or the DND1–NANOS3 complex identified by STREME (top row). (Bottom row) All motifs identified by STREME; y-axis indicates -log_10_ of p-value, percentage of binding sites containing each significantly enriched motif is indicated. **(d)** Empirical cumulative distribution function (CDF) of transcript abundance changes in HEK293 cells after expression of DND1–NANOS3 (left panel), DND1 alone (middle panel), or NANOS3 alone (right panel). The left schematic panel illustrates binning of transcripts according to whether they contained two or more N3-DREs (red line), one (orange line), or no N3-DRE in PAR-CLIP binding sites (light blue line), or no binding sites (black line). n = number of genes in each group; med.lfc = median of log₂ fold change in each group. **(e)** Same as in (d) but binning of transcripts according to whether the N3- DREs are in 3′ UTR (≥2 or 1), coding sequence (CDS), or 5′ UTR. Transcript abundance changes in HEK293 cells after expression of the DND1–NANOS3 (left panel), DND1 alone (middle panel), or NANOS3 alone (right panel). **(f)** Tolerance of the N3-DRE to single-nucleotide variation. All possible single-nucleotide variants of the N3-DRE were considered (top panel). Transcripts were binned according to the presence or absence of the N3-DRE variants in their 3′ UTR. Each tile on the bottom panel represents one N3-DRE variant, with the x-axis indicating the position in the motif and the y-axis indicating the substituted base. The color scale reflects the log_2_ of the median transcript abundance change of transcripts containing this N3-DRE variant upon DND1–NANOS3 expression.

Most mRNA binding sites for DND1, NANOS3, and DND1–NANOS3 were on exonic sequences (5′ untranslated region (UTR)/coding sequence (CDS)/3′ UTR) (Extended Data Fig. 2a), consistent with their predominantly cytoplasmic localization (Extended Data Fig. 1c). DND1 and the DND1–NANOS3 complex preferred 3′ UTRs for binding (Fig. 1b). In contrast, NANOS3 showed no preference for transcript regions, and its binding sites also lacked a specific sequence motif (Fig. 1b, c, Extended Data Fig. 2a), similar to a recent NANOS3 eCLIP in HEK293 cells^37^. As we and others previously reported^4,38^, DND1 alone bound a degenerate A- and U-rich motif (Fig. 1c, Extended Data Fig. 2b).

Remarkably, the complex of DND1 and NANOS3 bound at a sequence-defined heptamer, AUGAAUU (Fig. 1c). This motif, featuring a prominent G at the third position, cannot be explained as the composite of the individual sequence preferences of DND1 or NANOS3. While we did observe a subtle enrichment of AUG containing 5-mers in DND1 PAR-CLIPs (Extended Data Fig. 2b), this recognition is markedly enhanced by DND1-NANOS3 complex formation (Extended Data Fig. 2b), explaining its altered target mRNA preferences (Fig. 1a). Furthermore, more than 82% of mRNAs that were robustly expressed in HEK293 and contain the AUGAAUU motif in their 3′ UTR were bound by the DND1–NANOS3 RNP (Extended Data Fig. 2c), indicating that our Tandem fPAR-CLIP experiments exhaustively captured DND1–NANOS3 binding sites and that this motif is predictive and sufficient for DND1–NANOS3 binding. The AUGAAUU heptamer represents a *cis*-regulatory element potentially controlled by DND1–NANOS3, and thus, we termed it the NANOS3-dependent DND1 recognition element (N3-DRE).

### DND1–NANOS3 suppresses genes with a 3′-UTR N3-DRE

CCR4-NOT components co-purified in our tandem purification of the DND1–NANOS3 complex (Extended Data Fig. 1j), suggesting that DND1–NANOS3 represses target mRNAs through deadenylation, which is typically followed by decapping and mRNA degradation^39^. We therefore examined changes in target mRNA abundance upon induction of FH-DND1, NANOS3-VH, or both in HEK293 cells (Extended Data Fig. 2d). Expression of the DND1–NANOS3 complex suppressed mRNAs bound in the 3′ UTR with progressively stronger downregulation as the number of binding sites increased (median reduction: 3.8-fold for ≥7 sites and 2.3-fold for 4–6 sites, compared with 1–3 sites), whereas DND1 or NANOS3 alone did not appreciably suppress their targets (Extended Data Fig. 2e). mRNAs bound by the DND1–NANOS3 complex in CDS or 5′ UTR were much less affected by expression of the RNP (Extended Data Fig. 2f), indicating that binding in the 3′ UTR is necessary for DND1–NANOS3 to suppress target mRNAs, similar to other RBPs recruiting the CCR4-NOT complex, such as ZFP36 or the miRNA-induced silencing complex^40^.

We hypothesized that the N3-DRE was the *cis*-acting element preferentially controlled by the DND1–NANOS3 RNP. Indeed, the DND1–NANOS3 complex suppressed 3′ UTR targets containing the motif more strongly than those lacking the motif (Fig. 1d). The presence of the N3- DRE predicted mRNA suppression upon DND1–NANOS3 expression: 3,297 mRNAs contained the N3-DRE within their annotated 3′ UTRs, and of these, 2,770 (84%) were expressed in HEK293 cells (baseMean ≥ 10). Expressed mRNAs harboring even a single 3′-UTR N3-DRE were consistently and efficiently suppressed by the DND1–NANOS3 RNP, whereas mRNAs with a 5′ UTR or coding region N3-DRE were minimally affected (Fig. 1e).

To examine the N3-DRE sequence requirements, we systematically explored the tolerance of the N3-DRE to single-nucleotide substitutions. All single-nucleotide replacements, especially N3-DRE positions 1, 2, 3, and 7 (referred to as A1, U2, G3, and U7), attenuated DND1–NANOS3 RNP-mediated mRNA suppression in HEK293 cells, indicating sequence-dependent DND1–NANOS3 regulation (Fig. 1f).

We next examined whether the N3-DRE predicts DND1–NANOS3-mediated PTGR in germ cells. We induced murine PGCLCs^41^, profiled their transcriptome, and found that *Nanos3* KO results in robust upregulation of mRNAs containing N3-DREs in their 3′ UTR, whereas single-nucleotide-mismatch variants showed little or no effect (Extended Data Fig. 3a-c). We also reanalyzed RNA-seq data from FACS-sorted PGCs isolated from *Dnd1^-/-^* mice at E11.5^27^ or Dnd1 loss-of-function mutants (*Dnd1^ter/ter^*) at E12.5–14.5^38^, developmental timepoints at which Dnd1 and Nanos3 are co-expressed^2,14^. Dnd1 loss correlated with marked upregulation of mRNAs with 3′UTR N3-DREs (Extended Data Fig. 3d). These findings suggest that *in vivo* and in cultured germ cells, DND1 and NANOS3 cooperatively establish a gene expression profile critical for PGC survival through the N3-DRE. Collectively, our findings validate the N3-DRE in mRNA 3′ UTRs as a *cis*-element necessary and sufficient to predict DND1–NANOS3-mediated suppression.

### Cell cycle genes and epigenetic regulators contain N3-DREs and are downregulated by expression of the DND1–NANOS3 RNP

We noticed that co-induction of DND1 and NANOS3 for Tandem PAR-CLIP produced a pronounced reduction in cell proliferation, which may partially recapitulate cell cycle arrest and epigenetic reprogramming characteristic of the PGC migration phase^20,23^. We therefore systematically characterized the cell cycle and proliferation of HEK293 cells upon DND1 and NANOS3 induction. Induction of the DND1–NANOS3 RNP completely inhibited active cell proliferation within 24 h, whereas induction of either RBP alone had no effect (Fig. 2a). Cells co-expressing DND1 and NANOS3 were arrested at the G2/M and G1 phases (Fig. 2b). A comparable G2/M arrest was induced in mouse PGCLCs expressing DND1 and NANOS3 (Extended Data Fig. 3e). G2/M arrest in both germ and non-germ cell contexts indicates that this phenotype is intrinsic to DND1–NANOS3 activity.

**Figure 2.**
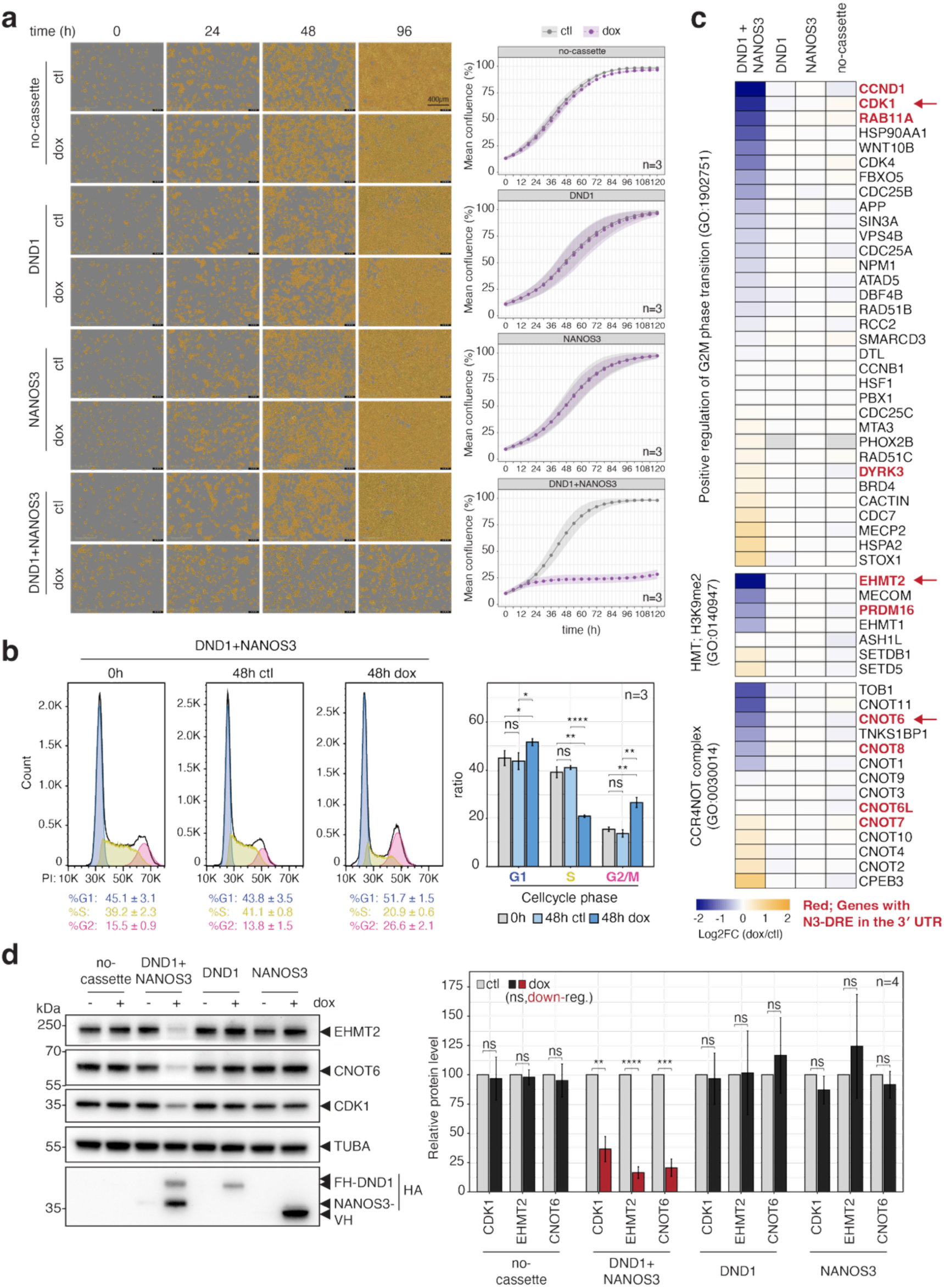
Co-expression of DND1 and NANOS3 induces cell cycle arrest along with the suppression of genes associated with cell cycle and reprogramming. **(a)** Effect of doxycycline (dox)-inducible expression of DND1, NANOS3, or DND1–NANOS3 on proliferation in HEK293 cells. Left: Brightfield images of cells following induction; cells are highlighted in yellow. Right: Growth curves with or without gene induction. Purple and gray shading indicate the standard deviation across time points. **(b)** Cell cycle distribution determined by propidium iodide-staining and flow cytometry before and 48 h after co-induction of DND1 and NANOS3. (Left) Cell cycle phases are shown in blue (G1), yellow (S), and pink (G2/M). (Right) Quantification of the proportion of cells in each phase. **(c)** Gene expression changes in pathways relevant to early PGC development. Expression changes were calculated as DESeq2 log₂ fold change in doxycycline-induced cells (dox) relative to non-induced controls (ctl). Top: positive regulation of G2/M transition (GO:1902751); Middle: H3K9me2 methyltransferase activity (GO:0140947); Bottom: CCR4-NOT complex (GO:0030014). Genes with the N3-DRE in the 3′ UTR are shown in red. Genes indicated by arrows were validated by WB. **(d)** Verification of protein expression for genes identified as downregulated in **c**. Left: Immunoblots of selected proteins. Right: Quantification of protein expression. Band intensities were normalized to TUBA within each sample, then further normalized to the dox/control. Error bars indicate standard deviation (S.D.) *\*p* < 0.05, *\*\*p* < 0.01, *\*\*\*p* < 0.001, *\*\*\*\*p* < 0.0001. n = number of independent experiments.

At a molecular level, the DND1–NANOS3 RNP likely binds directly and suppresses multiple transcripts encoding molecules involved in PGC-related processes. Indeed, 4 of 33 transcripts annotated as part of the positive regulation of G2/M phase transition (GO:1902751) contained canonical N3-DREs in their 3′ UTR (Fig. 2c) and were downregulated by DND1–NANOS3 expression, as were 2 of 7 genes that regulate methylation at histone H3 lysine 9 (GO:0140947) (Fig. 2c). In addition, we observed a reduction of co-immunoprecipitated CNOT6 in our Tandem PAR-CLIP preparation (Extended Data Fig. 1j) and found that it also contains the N3-DRE and showed reduced expression upon co-expression (Fig. 2c). We validated that reduced mRNA abundance upon DND1–NANOS3 complex expression corresponded to reduced protein levels, by assessing protein levels for a set of cell cycle and epigenetic regulators that contain the PAR-CLIP binding site with N3-DRE in their 3′ UTRs (EHMT2, CNOT6, CDK1) (Fig. 2d, Extended Data Fig. 4a-c). Collectively, we propose that the DND1–NANOS3 RNP regulates N3- DRE-containing transcripts, coordinating cell cycle arrest and chromatin remodeling, processes critical for PGC survival during migration-phase germ cell reprogramming.

### Structural basis for N3-DRE recognition by the DND1–NANOS3 complex

We determined a 1.7-Å resolution crystal structure that visualizes how DND1 and NANOS3 partner to recognize the N3-DRE (Extended Data Table 1). For crystallization, we used DND1 RRM1-RRM2 (referred to in structural and biochemical experiments as DND1), NANOS3 zinc finger (ZF) 1-ZF2 (NANOS3 ZF), and an RNA corresponding to the CDK1 N3-DRE (5′- auAUGAAUUu-3′) (Fig. 3a, Extended Data Fig. 4a). Together, DND1 and NANOS3 ZF formed a continuous RNA-binding surface that contacted all seven nucleotides of the N3-DRE (Fig. 3b). DND1 recognized the 5′-end of the N3-DRE, and NANOS3 recognized the 3′-end (Fig. 3c-i, Extended Data Fig. 5a-c). DND1 contacted nucleotides A1-A5 and formed hydrogen bonds that specifically recognized the A1 and U2 bases - also observed in a recent DND1 NMR structure^42^ - as well as G3, while NANOS3 contacted nucleotides A5-U7 and formed hydrogen bonds with bases U6 and U7. Stacking interactions with most bases added binding energy and could contribute to base specificity^43^.

**Figure 3.**
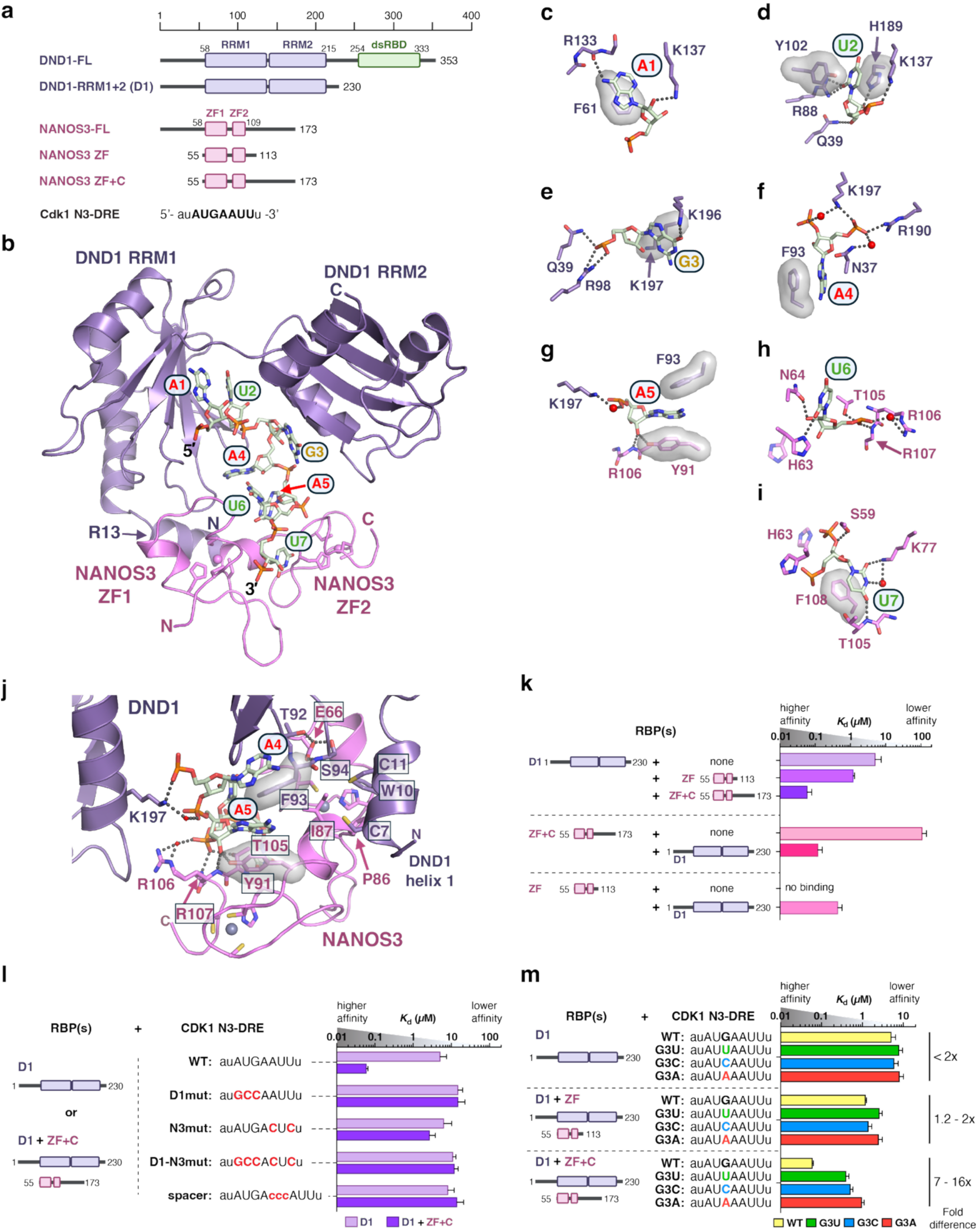
DND1 and NANOS3 collaboratively recognize the N3-DRE. (**a**) Schematic illustrations of human DND1 and NANOS3 protein domains. Protein constructs used in this study are indicated below. (**b**) Crystal structure of DND1 (purple) and NANOS3 (pink) reveals a continuous surface to recognize the 7-nt N3-DRE RNA. (**c-i**) Hydrogen bond, salt bridge, and stacking interactions formed between DND1–NANOS3 and N3-DRE nucleotides. Hydrogen bonds and salt bridges are indicated by dotted lines. Water molecules that mediate side chain- nucleotide interactions are represented by red oxygen spheres. Residues that stack with RNA bases are shown with transparent surfaces. (**j**) DND1 and NANOS3 recognize N3-DRE nt A5. Protein-protein and protein RNA interfaces near N3-DRE nts A4-A5 are highlighted. A primary DND1–NANOS3 interaction surface includes DND1 helix 1. (**k**) *In vitro* RNA-binding assays demonstrate that DND1 and NANOS3 ZF+C bind with high affinity to N3-DRE. RBP(s) included (left); *K*_d_ values measured by MST are shown as a bar graph (right). See also Supplementary Table 2. *K*_d_ values were determined from 3 independently pipetted measurements. (**l**) High-affinity RNA binding requires an intact N3-DRE sequence. Mutations of DND1 or NANOS3 binding sites (D1mut, N3mut, D1-N3mut) or separation of DND1 (A1-A4) and NANOS3 (A5-U7) interaction sequences by a spacer of three cytosines (spacer) abolishes high-affinity binding. (**m**) DND1–NANOS3 ZF+C selectively recognize G3. *K*_d_ values measured by MST are shown for binding to CDK1 N3-DRE RNAs with G3 (wt) or substituted with the other nts. Fold differences (right) indicate the range of differences relative to G3 (wt).

Notably, DND1 and NANOS3 both bound A5, and protein-RNA and protein-protein contacts in this region contributed to their joint RNA recognition. The A5 base was sandwiched between DND1 F93 and NANOS3 Y91 (Fig. 3g). DND1 K197 formed a water-mediated contact with the A5 phosphate group, and NANOS3 Y91 bound to its 2′OH. Multiple protein-protein interactions also structurally supported DND1 F93 and NANOS3 Y91 (Fig. 3j). NANOS3 E66 formed hydrogen bonds with DND1 T92 and S94, and the N-terminal helix of DND1 (C7, W10, C11) formed hydrophobic interactions with a short helix between the two NANOS3 zinc fingers (P86, I87). The specific RBP-RNA contacts thus provide a clear rationale for the motif specificity observed in cell-based assays (Fig. 1f).

### DND1 and NANOS3 partnership produces high-affinity binding to the N3-DRE

To expand upon the Tandem PAR-CLIP identification of the motif and crystal structure illustrating N3-DRE recognition, we analyzed the quantitative impact of the DND1–NANOS3 partnership on RNA-binding affinity (Supplementary Table 2). We measured collaborative binding to the CDK1 N3-DRE using microscale thermophoresis (MST) with DND1 (residues 1-230) and NANOS3 ZF (residues 55-113) or NANOS3 ZF+C (residues 55-173), which includes its C-terminal intrinsically disordered region (IDR) (Fig. 3a, Extended Data Fig. 5d-g).

Individually, DND1 (*K*_d_ = 5.1 μM) and NANOS3 ZF+C (*K*_d_ = 109 μM) bound to the CDK1 N3-DRE with modest affinity, and no RNA binding was detected for NANOS3 ZF (Fig. 3k). The presence of 2 μM NANOS3 ZF+C dramatically enhanced DND1 binding affinity 85-fold (*K*_d_ = 0.06 μM, Fig. 3k). NANOS3 ZF also improved DND1 affinity but only 4-fold (*K*_d_ = 1.2 μM). The NANOS3 C-terminal IDR is therefore critical for DND1–NANOS3 collaborative binding. Likewise, DND1 increased the RNA-binding affinity of NANOS3 (Fig. 3k), and DND1–NANOS3 ZF+C (*K*_d_ = 0.12 μM) bound with higher affinity than DND1–NANOS3 ZF (*K*_d_ = 0.44 μM). Together, our results demonstrate that the DND1–NANOS3 partnership not only creates a new recognition motif but is required for high-affinity binding.

Since DND1 interacted with nucleotides 1-5 of the N3-DRE, while NANOS3 interacted with nucleotides 5-7, we determined how the two half-sites contribute to overall binding affinity (Supplementary Table 2). When the DND1 half-site was mutated (D1mut: AUGA to GCCA), DND1 binding was disrupted (*K*_d_ = 15 μM), and NANOS3 did not enhance affinity (Fig. 3l). When the NANOS3 RNA site was mutated (N3mut: AUU to CUC), DND1 bound with modest affinity to its intact binding site (*K*_d_ = 6.4 μM), but NANOS3 did not increase affinity (Fig. 3l). Mutating both DND1 and NANOS3 sites (D1-N3mut) reduced RNA binding to levels seen with the individual mutants, with no signs of collaboration (Fig. 3l). Overall, these data indicate that high-affinity collaborative binding requires an intact N3-DRE with both DND1 and NANOS3 binding sites.

The continuous recognition surface for the N3-DRE with the central A5 bound by both proteins and our PAR-CLIP analyses suggested that the N3-DRE is a single motif, not a composite of the DND1 and NANOS3 preferences. We further confirmed this conclusion by inserting a 3-nt spacer (CCC) between the two half-sites, which eliminated high-affinity DND1–NANOS3 binding (Fig. 3l). Simultaneous recognition of A5 in the N3-DRE might be necessary for high-affinity interactions.

### Specificity for N3-DRE G3 depends upon the DND1–NANOS3 partnership

A striking revelation from the Tandem PAR-CLIP analyses was the specificity for a G at the third position of the N3-DRE (G3) not present in the DND1 motif (Fig. 1c). Our crystal structure indicates interactions between G3 and DND1 K196 and K197, but no interaction with NANOS3 (Fig. 3e). We measured the RNA-binding affinities of DND1–NANOS3 for CDK1 N3-DRE where G3 was substituted with the other nucleotides (Supplementary Table 2). DND1–NANOS3 ZF+C preferentially bound to G3 over U3, C3, or A3 (Fig. 3m). G3 specificity was not observed for DND1 alone or in combination with NANOS3 ZF (Fig. 3m). Therefore, G3 selectivity required the NANOS3 C-terminal IDR, which was absent from our crystal structure. We were unsuccessful in attempts to crystallize a complex with NANOS3 ZF+C. To identify the location of the NANOS3 C-terminal IDR (residues 114-173), we crosslinked DND1 and NANOS3 ZF+C bound to RNA with BS3, which covalently links nearby amine groups, such as lysine residues (Extended Data Fig. 5h). Mass spectrometry analysis identified intermolecular crosslinks among NANOS3 C-terminal IDR residues K130, K131, K139, and K160 and DND1 K196, K197, and K221, indicating that the NANOS3 C-terminus is proximate to K196/K197 that bind G3 (Extended Data Fig. 5i-j, Supplementary Table 3). Of note, we found that DND1 and NANOS3 ZF+C did not crosslink in the absence of RNA, suggesting that RNA drives the complex assembly (Extended Data Fig. 5h). This is consistent with a previous study where a DND1 mutant, deficient in RNA binding, lacked interaction with NANOS2^7^.

### RNA contacts by DND1 and NANOS3 are essential for the integrity of the ternary complex

We identified seven residues in DND1 and four in NANOS3 that contact RNA bases through their side chains (Fig. 4a). We assessed the relevance of each of these 11 residues by determining the effect of alanine substitution in our DND1–NANOS3 co-expression HEK293 cell line (Fig. 4b). Consistent with the importance of RNA binding for complex formation, all mutations abolished detectable DND1–NANOS3 co-IP (Extended Data Fig. 6a). We quantified the functional consequences of these mutations by measuring: (1) cell growth delay, (2) mRNA stability of N3- DRE containing genes using RNA-seq, and (3) target gene protein expression by Western blot (Fig. 4c). All DND1 (7 of 7) and two NANOS3 (2 of 4; Y91A and F108A) alanine substitutions lost the ability to arrest proliferation and suppress N3-DRE genes that we observed for WT proteins (Fig. 4d-e, Extended Data Fig. 6b-d). Substituting alanine for DND1 K196, which contacts G3, failed to repress cell proliferation and suppress N3-DRE genes (Fig. 4d), yet DND1 K196A retained *in vitro* RNA-binding activity (*K*_d_ = 0.08 μM vs 0.06 μM for WT). However, DND1 K196A lost selectivity for N3-DRE G3 *in vitro* (Extended Data Fig. 6e), indicating the importance of K196 for G3 selectivity and cellular function. Two NANOS3 mutations, H63A and K77A, did not affect the function of the DND1–NANOS3 RNP (Fig. 4e), although they had lost detectable interaction with DND1 by co-IP (Extended Data Fig. 6a).

**Figure 4.**
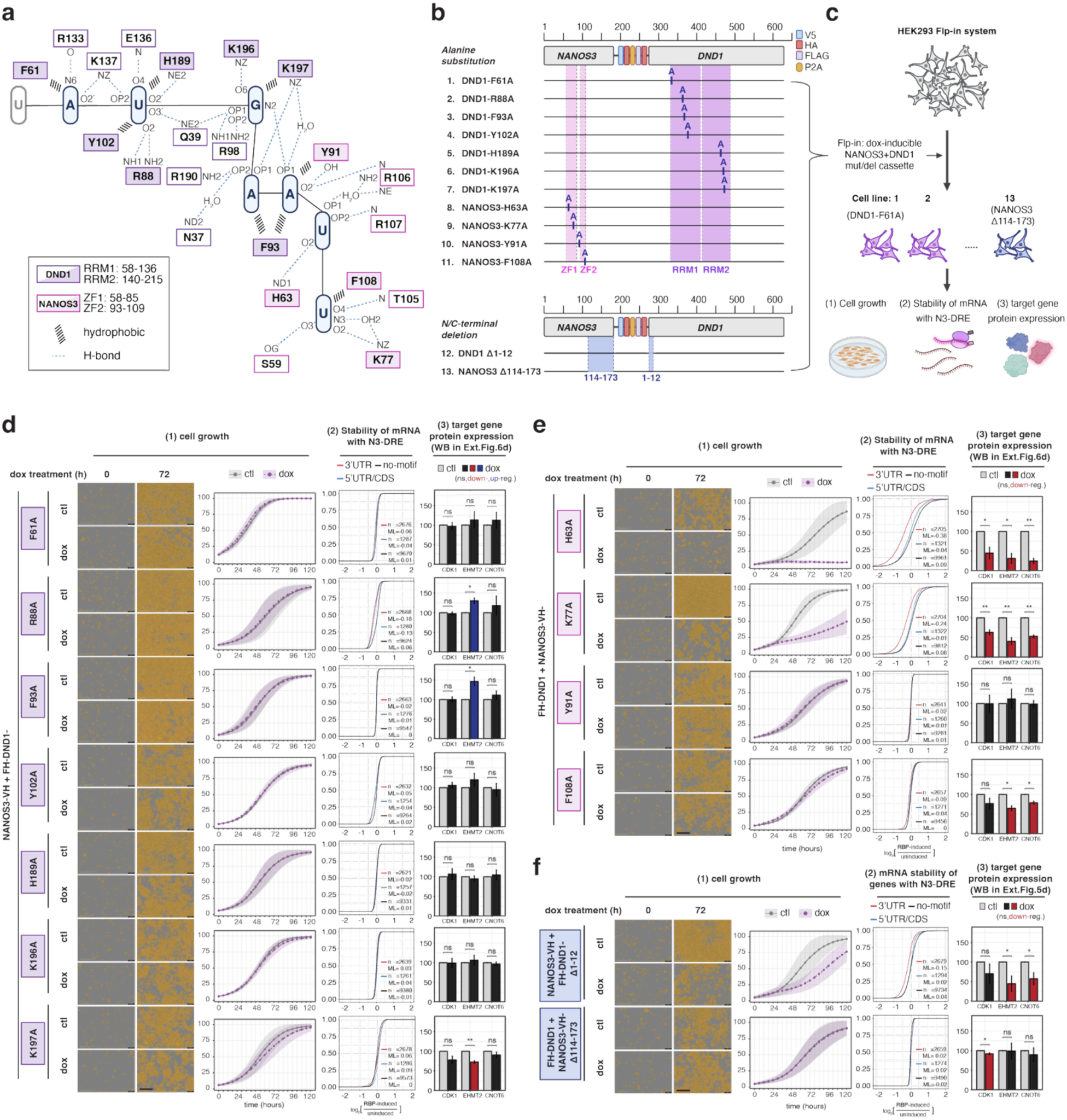
The RNA-binding residues in DND1 and NANOS3 and the C-terminal IDR of NANOS3 are essential for the integrity and function of the ternary complex. **(a)** Amino acid residues of DND1 (purple) and NANOS3 (pink) interacting with the N3-DRE. Residues with filled shading contact RNA bases through their side chains and were substituted with alanine to assess their importance for DND1–NANOS3 function. **(b)** Schematic of DND1–NANOS3 co-expression construct with mutated amino acids indicated (top) or DND1 and NANOS3 truncations highlighted in blue (bottom). **(c)** Schematic of assays performed with cells expressing constructs from (b). **(d)** Results from cell-based assays for DND1–NANOS3 constructs expressing DND1 point mutants. Left panels show results from IncuCyte growth assays. The middle panels show the effect of expression of mutant constructs on N3-DRE-containing RNAs. n = number of genes in each group; ML = med.lfc. Right panels show protein expression levels of three N3-DRE containing target genes (CDK1, EHMT2, and CNOT6) determined by Western Blot after expression of mutant constructs. The y-axis in each graph represents mean confluence, cumulative fraction, and relative protein level, respectively. **(e)** Same as in (d) except for the expression of DND1-NANOS3 constructs with point mutations in NANOS3. **(f)** Same as in (d) except for expression of DND1- NANOS3 constructs with truncations in DND1 or NANOS3. All experiments were performed in three independent biological replicates. *\*p* < 0.05, ** *p < 0.01*.

We also examined deletions of the N-terminal helix of DND1 (DND1 Δ1-12), the primary interaction surface with NANOS3 (Fig. 3j), and the C-terminal IDR of NANOS3 (NANOS3 Δ114- 173), which enhanced RNA-binding affinity of the DND1–NANOS3 RNP *in vitro* (Fig. 3k). In our cell culture model, DND1 Δ1-12 lost detectable association with NANOS3 but retained partial suppression of cell proliferation (Fig. 4f, Extended Data Fig. 6a), consistent with our *in vitro* data suggesting that it is not essential for RNA binding (*K*_d_ = 0.11 μM vs 0.06 μM for WT, Supplementary Table 2, lines 30-32). In contrast, and also in line with our *in vitro* RNA-binding data (Fig. 3k, NANOS3 ZF), NANOS3 Δ114-173 completely inactivated the DND1–NANOS3 RNP (Fig. 4f). We concluded that RNA recognition and binding specificity are functionally essential in living cells.

### Mutation of the N3-DRE attenuates DND1–NANOS3 effects in cultured cells and in vivo

To directly test whether the N3-DRE is sufficient to mediate repression of endogenous targets by the DND1–NANOS3 complex, we mutated the motif using CRISPR-Cas9 genome editing within an endogenous target gene. We selected *CDK1* because it is one of the top DND1–NANOS3 targets and harbors a single prominent and mammalian-conserved N3-DRE site bound by the complex (Fig. 5a, Extended Data Fig. 7a). In our HEK293 cell lines inducibly co-expressing DND1 and NANOS3, we mutated the *CDK1* N3-DRE from AUGAAUU to GCCACUC, a sequence that disrupts both the DND1 and NANOS3 binding sites and is bound poorly by the DND1–NANOS3 RNP (Fig. 5b, Extended Data Fig. 7b-e, Supplementary Table 2). RNA sequencing and Western blotting showed that disruption of the CDK1 N3-DRE specifically de-repressed CDK1 expression at both 24 and 48 h after induction of DND1 and NANOS3, without affecting other targets (Fig. 5c-e).

**Figure 5.**
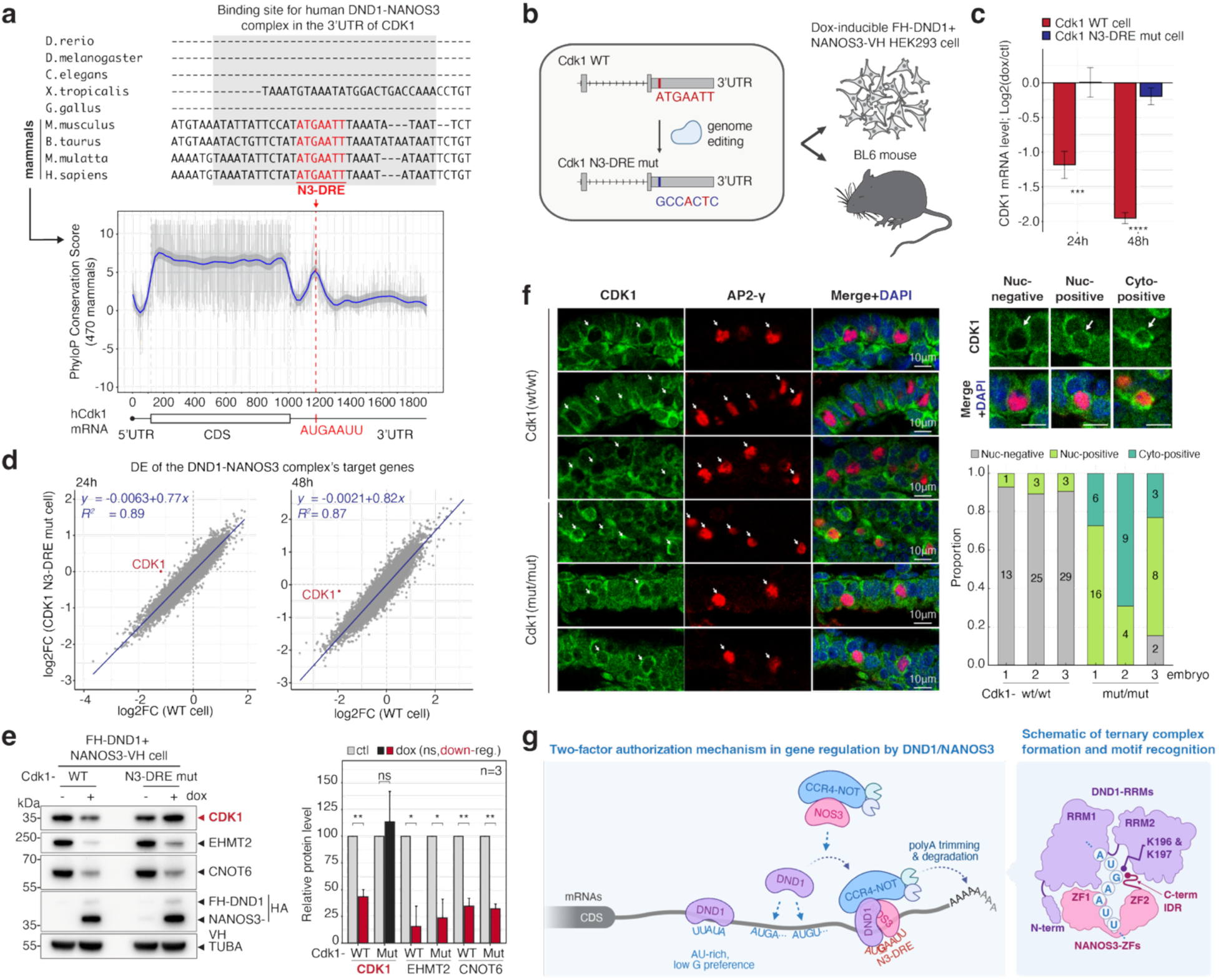
*CDK1* contains a mammalian-conserved N3-DRE essential for suppression. **(a)**. (Top panel) Conservation of the N3-DRE in *CDK1* gene across model organisms. 3′ UTR sequences from selected species were aligned using CLUSTAL Omega (v1.2.4). **(Bottom)** PhyloP conservation score from alignment of 470 mammalian genomes across the *CDK1* gene, showing that the N3-DRE is deeply conserved. Human *CDK1* mRNA (NM001786.5) defines 5′ UTR, CDS, and 3′ UTR positions. **(b)** Schematic of mutation strategy to change the N3-DRE at the *CDK1* locus in HEK293 cells and in mice. **(c)** *CDK1* mRNA levels in WT and *CDK1*-mutant cells 24 and 48 h after DND1–NANOS3 induction. **(d)** Scatter plot of DESeq2 log₂ fold changes for 3′ UTR target genes of the DND1–NANOS3 complex, comparing WT and *CDK1*-mut cells at 24 h and 48 h after DND1 and NANOS3 induction. Linear regression was performed, and regression coefficients (*r*) are shown in each plot. **(e)** Protein levels of three N3-DRE containing target genes (CDK1, EHMT2, and CNOT6) in WT and *CDK1*-mut cells at 48h after DND1–NANOS3 induction (left). Quantification of gene expression level (right). **(f)** (left) Representative IF images from WT and *Cdk1*-mut mouse embryos (E8.5). Cdk1 staining is shown in green, PGCs are marked in red by TFAP2C (AP2-gamma) stain, and nuclei are shown in blue after DAPI staining. Scale bar represents 10 μm. White arrows indicate AP2-gamma positive PGCs. (right) PGCs were manually classified into three Cdk1 distribution patterns (Nuc-negative, Nuc-positive, Cyto- positive). Counts were obtained across three sections per embryo, and per-embryo proportions were calculated and summarized in a bar graph. All PGCs within each section that had a clearly identifiable nucleus and proper focus were counted. **(g)** Motif recognition and gene regulation by the DND1–NANOS3 complex. (Left) Two-factor authorization mechanism in gene regulation by DND1 and NANOS3. DND1 primarily binds AU-rich sequences but can also weakly recognize AUG-containing elements. NANOS3 has weak RNA-binding capacity and no specific motif preference but interacts strongly with the CCR4-NOT complex. When DND1 and NANOS3 form a complex, they acquire stable and specific binding to the AUGAAUU (N3-DRE), resulting in selective suppression of N3-DRE-bearing genes in the 3′ UTR. (Right) Model of motif recognition by DND1 and NANOS3. The RRMs of DND1 bind to AUGAA, while the ZFs of NANOS3 bind to AUU, with both factors sharing recognition of the fifth A. The C-terminal IDR of NANOS3 is essential for RNA binding and gene suppression, while the N-terminal region of DND1 interacts with the ZF1 of NANOS3, contributing partially to RNA-binding activity but not being essential. Specificity for the third G in the N3-DRE is conferred by the C-terminal IDR of NANOS3 together with K196/197 in RRM2 of DND1. Data represent means from three biological replicates, with error bars indicating S.D. *\*p* < 0.05, ** *p < 0.01*, *\*\*\*p* < 0.001, *\*\*\*\*p* < 0.0001.

We next tested whether mutation of the *Cdk1* N3-DRE in mice would also disrupt control of Cdk1 levels in embryonic PGCs. We introduced the analogous mutation as above (Extended Data Fig. 8a, b) into the *Cdk1* N3-DRE in C57/BL6 mice using CRISPR-Cas9 genome editing (Extended Data Fig. 8c, d). Offspring from heterozygous intercrosses were distributed according to Mendelian ratios, indicating no embryonic lethality in the B6 background (Extended Data Fig. 8e). Immunofluorescence analysis of E8.5 embryos revealed that Cdk1 expression was suppressed in most TFAP2C (AP2-gamma)-positive PGCs in WT embryos (67/74, 90.5%), whereas nearly all PGCs in homozygous N3-DRE-mutant embryos exhibited Cdk1 de-repression (46/48, 95.8%) (Fig. 5f). Taken together, these results demonstrate that the N3-DRE alone mediates DND1–NANOS3-dependent repression in cultured cells and the embryonic germline.

## Discussion

Here, we identified the N3-DRE, a *cis*-acting element of high information content, found within target mRNA 3′ UTRs, and demonstrated that N3-DRE-dependent PTGR controls key processes required for PGC migration and development in the mammalian embryo. A detailed view of the molecular mechanisms underlying recognition of the N3-DRE by a heterodimeric complex comprising two RBPs, DND1 and NANOS3, illustrates the necessity of collaboration for sequence recognition and binding affinity (Fig. 5g). RNA binding is mediated not only by structured RNA-binding domains, but also the C-terminal IDR of NANOS3, which is essential for the stability and function of the complex.

The most notable feature of the N3-DRE is the emergence of a unique motif that is not merely an additive motif formed by the intrinsic specificities of both DND1 and NANOS3. Rather, the extended RNA interaction surface and specific contacts created by the recognition of two RBPs with low (DND1) and no (NANOS3) sequence specificity generate an unusually high-information-content regulatory element. Individual RNA-binding domains, such as the RRMs of DND1 and ZFs of NANOS3, typically recognize short (4-5 nt) stretches of sequence, even in tandem, and some positions in a longer motif may be degenerate. Remarkably, DND1–NANOS3 specifically interacts with each of the N3-DRE’s seven nucleotides, which are important for regulation (Fig. 1f). In its length and specificity, the N3-DRE resembles functional miRNA sites, which are 7-8 nts long, as well as 8-nt binding sites for Pumilio proteins^36,44-46^. The N3-DRE also strongly resembles a motif that was previously suggested as a preferred binding site for NANOS2 in murine SSCs^47^, AU(G/U)AA(A/U)UR. Considering that we and others^37^ did not identify sequence-specific RNA binding in HEK293 cells for either NANOS2 or NANOS3 alone (Fig. 1c) and that DND1 is expressed in SSCs, we propose that the study^47^ captured DND1–NANOS2 binding sites^6,7^.

The N3-DRE controls a powerful gene expression circuit critical for fertility, and its aberrant activation in any cell type will result in severe phenotypes. Splitting the effector complex into two parts—DND1 and NANOS3—that individually cannot recognize the N3-DRE and tamper with the gene regulatory circuit serves as a form of two-factor authorization protecting cells from aberrant expression of individual components. Such a mechanism exists at the transcriptional level; for example, two-factor authorization protects the germline by silencing transposable elements^48^ through an essential partnership of two chromatin readers, SPIN1 and the piRNA-induced silencing complex-recruiting protein SPOCD1.

The DND1–NANOS3 RNP may serve as a paradigmatic example for collaborative RNA recognition. RBPs are often found in larger heterotypic RNPs; however, whether these larger RNPs recognize sites that are very different from the sum of their subcomponents is unclear. Most approaches for characterization of RBP specificity *in vitro* (RBNS^11^, SELEX^12^, RNAcompete^9^, etc.) or in cells (CLIP^10^, TRIBE^13^, etc.) study each RBP in isolation. A recent CLIP-based study identified proteins co-bound with selected RBPs^49^; however, without attempting to catalogue *cis*- acting elements, and a study of DND1 and NANOS3 function in human PGCLCs identified mRNAs bound by both proteins using iTRIBE but failed to identify individual or shared motifs^8^. *In vitro* selection has identified changes in RNA recognition sequences of *Drosophila melanogaster* and *Caenorhabditis elegans* PUF proteins with their partner proteins^50-52^. For the protein pairs studied, RNA-binding affinity increases vs. the PUF protein alone, yet the RNA sequence specificity is reduced by the partnerships. This is quite different from what we observed here for DND1–NANOS3, and future work geared towards identifying *cis*-acting elements will clarify how prevalent the phenomenon we reported here is, i.e., the generation of novel binding specificities by complex formation of two or more RBPs.

## Author contributions

E.F., M.B., A.J., M.S., Ma.Y, Mi.Y, and W.H. generated RBP-expressing HEK293 cell lines and conducted cell and molecular biology experiments. A.J. performed initial PAR-CLIP experiments that led to the discovery of CDK1 as the top target. M.S. performed PAR-CLIP and Tandem PAR-CLIP experiments, A.P. and M.S. analyzed them. M.S., A.J., Ma.Y., and Mi.Y. collected RNA-seq data, and M.S. analyzed them together with A.P. Ma.Y. and Mi.Y. generated mouse ESCs and conducted PGCLC induction assays. M.S. generated CDK1-N3-DRE mutant HEK293 cells. C.L. and M.S. generated CDK1-N3-DRE mutant mice, and M.S. performed animal experiments with the support of C.L. and X.F. C.Q. expressed recombinant DND1 and NANOS3 proteins and conducted X-ray crystal structural analyses of the DND1/NANOS3/RNA complex. C.Q. conducted MST RNA-binding experiments. C.Q. prepared crosslinked samples, and J.W. conducted mass spectrometry analysis and processed the data. M.S., M.B., and D.R. collected microscopy images. J.W., V.S., E.V., Ma.Y., M.H., and T.H. supervised experiments, analyzed data, and acquired funding. M.S., C.Q., and T.H. created and/or finalized all figures. M.S., C.Q., and T.H. wrote the initial draft of the manuscript and finalized it together with Ma.Y., T.H., and M.H.. All authors edited the manuscript.

## Acknowledgements

We thank the beamline staff for assistance with data collection at the National Synchrotron Light Source II beamline 17-ID-1 (AMX) and Lars Pedersen for crystallographic and data collection support at NIEHS. The authors acknowledge use of the NIEHS Structural Biology Core (ZIC ES102645) and NIEHS Mass Spectrometry Research Center (ZIC ES103005). We thank Dr. Stefania Dell’Orso, Faiza Naz, and Shamima Islam (NIAMS Genomics Technology Section) for high-throughput sequencing, and Aster Kanea (NIAMS Light Imaging Section) for help with acquiring microscopy images. We appreciate the help of James Simone (NIAMS Flow Cytometry Section) with flow cytometry. The authors acknowledge Dr. Rajagopal Chari (Genome Modification Core, CCR/NCI/NIH) for generating the Cas9 and donor plasmids used to establish the Cdk1 N3-DRE mutant cells. Figures partially created in BioRender. Suzawa, M. (2025) [ https://BioRender.com/n1llmw6]. This research was supported by the Intramural Research Program of the National Institutes of Health (NIH). The contributions of the NIH authors are considered Works of the United States Government. The findings and conclusions presented in this paper are those of the authors and do not necessarily reflect the views of the NIH or the U.S. Department of Health and Human Services.

## Material and Methods

### Cell lines and culture

Flp-In™ T-REx HEK293 cells (Thermo Fisher Scientific, R78007) were cultured in Dulbecco’s Modified Eagle Medium (DMEM; Gibco, 11995-065) supplemented with 10% fetal bovine serum (FBS; GeminiBio, 100-800-500), 1% penicillin-streptomycin (Gibco, 15140-122), and 15µg mL^-1^ blasticidin (Thermo Fisher Scientific, A1113902), at 37°C in a humidified incubator with 5% CO₂. HEK293 cell lines stably integrating a doxycycline (dox)-inducible cassette encoding epitope-tagged RBPs — including FLAG/HA (FH)-DND1, NANOS3-V5/HA (VH), NANOS3- VH-P2A-FH-DND1, and various DND1 and NANOS3 mutants or deletions — were generated by Flp recombinase-mediated integration using the Flp-In™ system, as previously described^53^. They were maintained in the same culture medium with the addition of 100 μg mL^-1^ Hygromycin B (Thermo Fisher Scientific, 10687010). Expression of tagged RBPs was induced by adding 2 μg mL^-1^ doxycycline (Millipore Sigma, D9891).

We previously^4^ established embryonic stem cells (ESCs) from the 129/Sv strain of mice (129S6/SvEvTac), and the same parental ESCs were maintained with N2B27/2iLIF medium supplied with 1% knockout serum replacement (KSR) (Thermo Fisher Scientific; cat. no. 10828- 010) on cell culture plates coated with 0.1% (w/v) gelatine as described previously^54^.

### CRISPR–Cas9-mediated gene editing for generating Nanos3 mutant ES cells

CRISPR–Cas9-mediated genome editing was performed as described previously^4^. We purchased a cocktail of three small guide RNAs (sgRNAs) (Santa Cruz Biotechnology, Inc.; cat# sc-434074), cloned individual sgRNA plasmids, and used pairs of the sgRNAs for the targeted deletions of the zinc-finger region of the *Nanos3* locus. The targeted deletion was confirmed by PCR genotyping. The PCR products were cloned into a pGEM-T Easy plasmid (Promega, cat. no. A1360), and approximately 10 clones per ES cell line were sequenced.

### Induction of mouse PGCLCs from ES cells

1.5 × 10^5^ cells per well of ES cells were plated on a 12-well plate and induced into EpiLCs for 48 h as described previously^4^. The EpiLCs were then cultured under a floating condition by plating 2–3 × 10^3^ cells per well of a 96-well Clear Round Bottom Ultra Low Attachment Microplate (Corning, cat. no. 7007) in GK15 medium [GMEM (SIGMA Aldrich; cat#G5154) containing 2 mM L-glutamine (Thermo Fisher Scientific; cat#11710-035) with 1 × MEM non-essential amino acids (Thermo Fisher Scientific; cat#11140-050), 1 mM sodium pyruvate (Thermo Fisher Scientific; cat#11360-070), 100 μM 2-mercaptoethanol (Thermo Fisher Scientific; cat#21985- 023), 1×10^3^ units/ml LIF (SIGMA-Aldrich; cat#ESG1107), and 15%(v/v) KSR] supplemented with LIF, BMP4 (500 ng ml^−1^), SCF (100 ng ml^−1^), and EGF (50 ng ml^−1^). Day 4 PGCLCs were purified with a fluorescence-activated cell sorter (FACS) (ARIA II; BD Biosciences) by using anti-SSEA1/CD71-eFluor660 (1:20 dilution; eBioscience, cat. no. 50-8813-41) and anti- ITGB3/CD61-PE antibodies (1:200 dilution; BioLegend, cat. no. 104307).

### Plasmid construction

pFRT/TO-FH-DND1, pFRT/TO-NANOS3-VH, and pFRT/TO-NANOS3-VH-P2A-FH-DND1 were generated as previously described^4^ Plasmid pFRT/TO human NANOS3 (hNOS3)-V5-HA (VH) or hNOS3-VH-P2A-FLAG-HA(FH)-hDND1 was generated by gBlock gene synthesis (Integrated DNA Technologies) of the respective coding sequences (CDS) followed by restriction digest with HindIII and EcoRV and ligation into pFRT/TO/FLAG/HA-DEST plasmid (Thermo Fisher Scientific) to create pFRT/TO hNOS3-VH or hNOS3-VH-P2A-FH-hDND1, respectively. Alanine substitutions at key RNA-binding residues, as well as N- and C-terminal deletion of DND1 and NANOS3, were introduced into pFRT/TO-NANOS3-VH-P2A-FH-DND1 using the Q5® Site-Directed Mutagenesis Kit (NEB, E0554S) according to the manufacturer’s instructions, with primers listed in Supplementary Table 4. All constructs were verified by Sanger sequencing.

### fPAR-CLIP

fPAR-CLIP was performed as previously described^35^ with modifications. For single fPAR-CLIP experiments using FH-DND1 and NANOS3-VH HEK293 cells, 2.0 × 10⁶ cells were seeded per 10 cm dish (1 to 3 dishes per replicate). Upon reaching ∼70% confluency, cells were treated with 2 μg mL^-1^ doxycycline for 24 h to induce expression of the RBPs. Cells were then labeled with 100 μM 4-thiouridine (4SU; Sigma-Aldrich, T4509) for 16 h, followed by UV crosslinking at 312 nm (0.5 J/cm²). Crosslinked cells were lysed in NP-40 lysis buffer (20 mM Tris-HCl pH 7.5, 150 mM NaCl, 2 mM EDTA, 1% NP-40, 0.5 mM DTT) supplemented with protease inhibitor cocktail (cOmplete™, Sigma-Aldrich, 4693159001) and phosphatase inhibitor cocktail (PhosSTOP, Sigma-Aldrich, 4906837001), and incubated on ice for 10 min. Lysates were cleared by centrifugation and treated with RNase T1 (1 U μl^-1^) for 15 min at 22°C. IP was performed using RNase-treated lysates incubated with anti-FLAG M2 magnetic beads (40 μl slurry per mL lysate; Sigma-Aldrich, M8823) for FH-DND1, or anti-V5 magnetic beads (300 μl slurry per mL lysate; MBL, M167-11) for NANOS3-VH, for two h at 4°C with rotation. Beads were washed three times with NP-40 lysis buffer and three times with high-salt wash buffer (identical to NP-40 lysis buffer but containing 500 mM NaCl). Subsequent steps — including dephosphorylation, on-bead 3′ adapter ligation, SDS-PAGE, excision of RNP–adapter bands, and RNA purification — were performed according to the standard fPAR-CLIP protocol^35^.

Tandem fPAR-CLIP was performed to isolate RNAs bound by the FH-DND1–NANOS3- VH complex. First, NANOS3-VH-P2A-FH-DND1 HEK293 cells were seeded using either eight 10 cm dishes (2.0 × 10⁶ cells per dish) or five 15 cm dishes with an equivalent total number of cells per replicate. Cells were treated with doxycycline, labeled with 4SU, and UV-crosslinked as described above. Following lysis and RNase T1 digestion, the first IP was performed using the V5-Tagged Protein Purification Kit Ver. 2 (MBL, 3317) according to the manufacturer’s instructions. Lysates were incubated with V5-agarose beads to capture NANOS3-VH and associated FH-DND1–NANOS3-VH complexes, washed with the kit-supplied buffer, and eluted using the included V5 peptide. The eluate was then subjected to a second IP using anti-FLAG M2 magnetic beads (40 μl slurry per ml of eluate; Sigma-Aldrich, M8823) to enrich for the FH-DND1–NANOS3-VH complex. Beads were washed as described for single fPAR-CLIP. All subsequent steps — including dephosphorylation, 3′ adapter ligation, SDS-PAGE, excision of RNP–adaptor complexes, and RNA purification — were carried out following the standard fPAR-CLIP workflow^35^. All cDNA libraries were prepared from purified RNA and sequenced as 50 bp single-end reads on an Illumina NextSeq 550, NextSeq 2000, or NovaSeq 6000 platform, and data were analyzed as follows.

PAR-CLIP data analysis was carried out using PCLIPtools (v0.7.2) (https://github.com/paulahsan/pcliptools). FASTQ data for the PAR-CLIP experiment were aligned to the respective genome assembly by the ‘pcliptools-align’ script of PCLIPtools. ‘pcliptools-align’ collapses the FASTQ to FASTA, removes duplicates, and removes the 5′ and 3′ adapters by using cutadapt (v4.5). The resulting HEK293 cell line-derived FASTA files were aligned to the hg38 genome, and mouse PGCLC*-*derived FASTA files were aligned to the mm10 genome with the STAR aligner (v2.7.9a). Since adapters were removed while generating FASTA files and short read alignments were expected, an ‘end-to-end’ alignment mode was set for STAR. For the alignment, the maximum number of mismatches was set to 2. Additionally, ‘–outFilterMultimapNmax’ was set to 10 to limit the maximum number of multi-mappers for a read to 10.

After generating a Binary Alignment Map (BAM) file, the ‘pcliptools-cluster’ script from PCLIPtools was used to call peaks. PCLIPtools calls peaks by using non-T-to-C mismatch information to compute the background mismatch rate and fits the signal into a Poisson distribution. Regions with higher observed T-to-C compared to the expected T-to-C are high-confidence peaks for the PAR-CLIP data that indicate potential interaction sites (clusters) for RNA and RBPs. An ENSEMBL^55^-generated Gene Transfer Format (GTF) file was used to annotate the clusters to respective genomic regions such as 5′ UTRs, coding sequences (CDS), 3′ UTRs, introns, miRNA etc. Minimum read depth for clusters was set to 3 reads. Clusters were further filtered by read depth and number of unique locations with T-to-C mismatch (loc_T2C), and then ranked by count of T2C in each cluster (count_T2C).

### *5-mer* analysis

An in-house script was used to compute the relative abundance of 5-mers in the PAR-CLIP cluster regions that depict over-represented sequences in RBP-RNA interaction sites. For our analysis, 200 nt regions upstream and downstream of the clusters were considered as background. The script utilizes BEDtools (v2.31.1) to extract FASTA sequences for both cluster regions and background regions. Afterward, the script computes a count of all possible 5-mers in both sets using Jellyfish^56^. Then the script calculates Z-score enrichment by proportion; 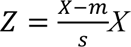 is the proportion of a given 5-mer in background regions. Additionally, μ the mean and standard deviation of the respective 5-mer for the background.

### Motif analysis

Motif enrichment analysis was performed using STREME v5.5.2^57^. Enriched 5- to 8-mer motifs were identified from the top 2,000 clusters upon filtering by minimum of 10 reads per cluster, ranked by the number of crosslinked events, obtained from both single and Tandem PAR-CLIP experiments. Motif discovery was terminated when three consecutive motifs exceeded the p-value threshold of 0.05. All identified motifs are shown in Fig.1c.

### IncuCyte cell proliferation assay

Cell proliferation was monitored using the IncuCyte® S3 Live-Cell Analysis System (Sartorius) based on confluence measurements over time. The HEK293 cell lines were seeded in 24-well plates at a density of 0.4 × 10^5^ cells per well, in triplicate for each condition. After cell attachment (∼24 h), 2 μg mL^-1^ doxycycline (or vehicle control) was added to induce RBP expression. Plates were placed into the IncuCyte system housed in a humidified 37°C incubator with 5% CO₂. Nine images per well were acquired every 6 h for up to 120 h using the 10× objective lens. Cell confluence (%) was calculated using the IncuCyte image analysis software (v2022B Rev2) with default settings, except for applying a 100 μm² area filter to exclude small debris during confluence calculation. Mean confluence values from triplicate wells were treated as a single biological replicate. The mean and standard deviation from three independent biological replicates were plotted over time to assess cell proliferation under each condition.

### Immunoblotting and antibodies

To assess protein expression, the HEK293 cell lines were seeded in 6-well plates at a density of 0.3 × 10^6^ cells per well. The following day, cells were treated with doxycycline (2 μg mL^-1^) and incubated for 48 h. After induction, 3 × 10⁶ cells per sample were harvested and lysed on ice for 10 min in 500 μl NP-40 lysis buffer supplemented with protease and phosphatase inhibitor cocktails. Lysates were cleared by centrifugation, and proteins were resolved by SDS-PAGE and transferred to PVDF membranes (Bio-Rad, 1704157). Membranes were blocked with 5% (w/v) skim milk in TBST (TBS + 0.1% Tween-20) for one h at room temperature, incubated overnight at 4°C with primary antibodies, and then with HRP-conjugated secondary antibodies for 1 h at room temperature. Signals were detected using SuperSignal West Pico PLUS Chemiluminescent Substrate (Thermo Fisher Scientific, 34580) and imaged with a ChemiDoc XRS+ system (Bio-Rad). Signal intensities were quantified using Image Lab 6.1 Software. Primary and secondary antibodies are listed in Supplementary Table 4.

### Conservation analysis across mammalian Cdk1 transcript

Conservation analyses were performed using the UCSC Table Browser (https://genome.ucsc.edu/cgi-bin/hgTables)^58^. The following settings were used: Clade – Mammal, Genome – Human, Assembly – Dec. 2013 (GRCh38/hg38), Group – Comparative Genomics, Track – Hiller Lab 470 Mammals, Table – 470 phyloP (phyloP470BW), Position – chr10:60778478–60794852. The retrieved phyloP scores were plotted according to the human CDK1 transcript (NM_001786.5).

### RNA-sequencing

RNAs from series of HEK293 cell lines at designated time points (0 to 48 h) were isolated with the Direct-zol RNA Miniprep Kit (Zymo Research, Cat# R2050) according to the manufacturer’s instructions. 500ng of total RNA was subjected to PolyA selection, then converted into a cDNA library and sequenced on an Illumina NovaSeq 6000 or NovaSeq X plus platform (paired-end, 50 or 100 bp). Reads from HEK293 cell lines and mouse PGCLCs were aligned to the hg38 and mm10 genome, respectively, using STAR (v2.7.9a)^59^. Multimappers were handled with parameter ‘ --outFilterMultimapNmax 20’ by allowing a maximum 20 multiple alignments for a read. The BAM file generated by STAR was used to calculate the raw counts for the genes by using featureCounts (v2.0.3)^60^. Genome-specific (i.e., hg38 and mm10) Gene Transfer Format (GTF) generated by UCSC RefSeq (https://hgdownload.soe.ucsc.edu/downloads.html) were used as gene annotation for featureCounts. To identify differentially expressed genes, DESeq2 (v1.36.0)^61^ was used in an R (v4.2.0)^62^ environment. DESeq2 normalized the raw count data to account for the depth of the libraries and the respective experimental factors, i.e., genotypes, and then generated normalized counts. Using the normalized counts it also estimated fold changes for genes across conditions, and the respective statistical confidence was also generated. These metrics were used to identify differentially expressed genes. Genes with adjusted p-value ≤ 0.05 and |log_₂_FC| > 1 were considered significant. Gene sets related to primordial germ cell (PGC) processes were obtained from AmiGO2^63-65^, specifically: “positive regulation of G2M phase transition” (GO:1902751), “histone H3K9me2 methyltransferase activity” (GO:0140947), and “CCR4-NOT complex” (GO:0030014). Previously published RNA-seq datasets profiling the PGC transcriptome from Dnd1⁻/⁻ mice at E11.5^27^ and Dnd1Ter/Ter mutants at E12.5–E14^38^ were retrieved from the GEO database under accession numbers GSE218770 and GSE132719, respectively.

Full DESeq2 outputs are provided in Supplementary Tables 5-8: Supplementary Table 5 (DND1, NANOS3, and DND1 and NANOS3 co-expression in HEK293 cells), Supplementary Table 6 (DND1 and NANOS3 point-mutant/deletion analyses), Supplementary Table 7 (*CDK1* N3-DRE mutant datasets), and Supplementary Table 8 (WT and *Nanos3*-KO mPGCLC datasets).

### Flow cytometry

The HEK293 cell lines were seeded at a density of 1.0 × 10⁶ cells per 10 cm dish. When cells reached approximately 20% confluency, samples designated as 0 h were collected, and the remaining dishes were treated with either 2 μg mL^-1^ doxycycline or vehicle control for 48 h. For each time point, 2 × 10⁶ cells were harvested, washed once with PBS, and pelleted by centrifugation. The cell pellet was resuspended in 200 μl PBS, and 500 μl of 100% ice-cold ethanol was added dropwise while gently mixing to fix the cells. Samples were kept on ice for 15 min, then centrifuged at 400 × g for 8 min at 4°C to remove the fixation solution. Cells were washed once with PBS and resuspended in 150 μl of propidium iodide (PI)/RNase staining solution (PBS containing 2 μg mL^-1^ PI and 100 μg mL^-1^ RNase A). Samples were incubated overnight at 4°C. Flow cytometry was performed using a FACS Celesta instrument (BD Biosciences), and data were analyzed with FlowJo software (v10.10.0, BD Biosciences). Single cells were first gated using forward scatter (FSC) versus side scatter (SSC) to exclude debris, followed by PI-A versus PI-W gating to exclude doublets (Supplementary Fig. 1). Cell cycle distribution (G0/G1, S, G2/M phases) was determined based on DNA content profiles.

### Generation of Cdk1 N3-DRE mutant HEK293 cells

Guide RNAs were designed using sgRNA Scorer 2.0^66^ with sequence spanning the specific region in the 3′ UTR as input sequence and subsequently tested for cutting activity in 293Ts cells stably expressing Cas9 using methodology previously described^67^. The guide RNAs designed and tested are listed in Supplementary Table 4. Candidates 3755 and 3759 were the most effective as measured by indel frequency and subsequently used for targeting experiments. Oligonucleotides corresponding to these two guides were annealed and phosphorylated and ligated into the pDG458 backbone^68^ (Addgene #100900) to generate the plasmid pCE1034. pDG458 was a gift from Paul Thomas (Addgene plasmid # 100900; http://n2t.net/addgene:100900; RRID: Addgene100900). To generate the donor carrying the 3′ UTR cluster mutations, DNA carrying the mutation along with regions of 5′ and 3′ homology was synthesized (Twist Biosciences) and cloned into the pGMC00018 (Addgene #195320) backbone using isothermal assembly^67^. pGMC00018 (aka pRC0082) was a gift from Raj Chari (Addgene plasmid # 195320; http://n2t.net/addgene:195320; RRID:Addgene 195320). All constructs were verified using Sanger sequencing.

NANOS3-VH-P2A-FH-DND1 HEK293 cells were seeded in 10 cm dishes at a density of 4.0 × 10⁶ cells per dish. The following day, cells were transfected with pCE1034 (SpCas9-2A-GFP and sgRNAs) and the donor plasmid using Lipofectamine 2000 (Thermo Fisher Scientific), according to the manufacturer’s instructions. At 48 h post-transfection, GFP expression was confirmed by fluorescence microscopy as an indicator of transfection efficiency. Subsequently, 6 × 10⁶ cells were resuspended in 1 mL PBS and subjected to fluorescence-activated cell sorting (FACS) using a FACSAria Fusion cell sorter (BD Biosciences). Cells with high GFP intensity (Top2%) were single-cell sorted into five 96-well plates (one cell per well). Colony formation was monitored over time, and wells containing single, isolated colonies were expanded sequentially to 24-well and then 12-well plates. Once colonies reached sufficient confluency, approximately 2.5 × 10⁶ cells were harvested, and genomic DNA was extracted using the QIAamp DNA Mini Kit (QIAGEN, 51304) according to the manufacturer’s protocol. The targeted genomic region was PCR-amplified and analyzed by Sanger sequencing to validate successful editing. The PCR primers used for genotyping are listed in the Supplementary Table 4.

### Protein expression and purification

cDNAs encoding DND1 constructs (DND1, residues 1-230; DND1 Δ1-12, residues 13-230) and NANOS3 constructs (ZF, residues 55-113; ZF+C, residues 55-173) were each subcloned into the pSMT3 vector with an N-terminal His_6_-SUMO tag. The plasmids were transformed into *E. coli* BL21-CodonPlus (DE3)-RIL competent cells (Agilent). A 1-L Terrific Broth culture supplemented with 50 µg mL^-1^ kanamycin was inoculated with a 10-mL overnight culture and grown at 37°C to an OD_600_ of ∼0.8. The media for NANOS3 also included an additional 0.1 mM ZnSO_4_. Protein expression was induced with 0.1 mM IPTG at 16°C, and the culture was grown overnight.

DND1 and NANOS3 proteins were purified following a similar protocol. The *E. coli* cell pellet was resuspended in a buffer containing 20 mM Tris pH 8.0, 0.5 M NaCl, 20 mM imidazole, 5% (v/v) glycerol, and 1 mM DTT and sonicated to lyse cells. Clarified lysate was mixed with 5 mL Ni-NTA resin (Qiagen) on a rotary platform for 1 h. The protein was eluted off Ni-NTA resin in a gravity column with a buffer containing 20 mM Tris (pH 8.0), 0.05 M NaCl, 0.5 M imidazole, and 1 mM DTT after extensive washing with the lysis buffer. The Ni-NTA eluate was incubated with Ulp1 protease at 4°C for 2 h to remove the His_6_-SUMO tag. Subsequently, the protein solution was filtered and loaded onto a Hi-Trap Heparin column (Cytiva). Heparin column buffer A contained 20 mM Tris (pH 8.0) and 1 mM DTT, and buffer B contained an additional 2 M NaCl. The column was eluted with a gradient of increasing buffer B from 5% to 100%. Heparin column peak fractions were concentrated to 5 mL and loaded onto a HiLoad 16/60 Superdex 75 column (Cytiva) in a buffer containing 20 mM HEPES pH 7.4, 150 mM NaCl, and 1 mM DTT. Purified protein was concentrated, flash frozen in liquid nitrogen, and stored at -80°C. Protein concentration was determined using A_280_ measured by NanoDrop and the calculated extinction coefficients of the proteins.

### Crystallization and structure determination

Purified DND1 1-230 protein, NANOS3 ZF protein and a 10-nt *CDK1*-N3-DRE RNA (5′- auAUGAAUUu-3′) were mixed at a molar ratio of 1:1.1:1.2 and incubated at 4°C for at least 1 h. The DND1/NANOS3/RNA ternary complex was purified with a Superdex 75 Increase 10/300 GL column (Cytiva) in a buffer containing 20 mM HEPES pH 7.4, 150 mM NaCl, and 1 mM DTT. The complex was concentrated to A_280_ ∼ 5, A_260_/A_280_ = 1.4. Crystals of the complex were grown by hanging drop vapor diffusion at 20°C with 2 µl sample + 2 µl reservoir solution. Microseeding was performed during crystallization optimization. Crystals were cryoprotected using crystallization solution supplemented with 25% (v/v) glycerol and flash frozen in liquid nitrogen.

X-ray diffraction data were collected at 100 K at beamline 17-ID-1 (AMX) of the National Synchrotron Light Source II at a wavelength of 0.92 Å. Data were autoprocessed using autoProc^69^. A criterion of CC1/2 > 0.3 was used for the resolution cutoff. The crystal belonged to the P2_1_ space group. An asymmetric unit contained one ternary complex. A structure model of the complex was first generated using AlphaFold^70^. The structure was divided into three individual models: DND1 RRM1, DND1 RRM2, and NANOS3 ZF, which were used for molecular replacement with Phaser^71^. Initial refinement revealed clear density for the RNA. The model was improved through iterative refinement and manual building with Phenix^72^ and Coot^73^. The final model contains DND1 residues 4-229, NANOS3 residues 55-112, and RNA nucleotides 3-9. 98.6% of protein residues are Ramachandran favored with no outliers. Data collection and refinement statistics are shown in Extended Data Table 1.

### Microscale Thermophoresis (MST) RNA-binding assay

5′-Cy5-labeled *CDK1*-N3-DRE-based wild-type and mutant RNAs were ordered from Horizon Dharmacon. To measure DND1-RNA dissociation constants, DND1 protein was two-fold serially diluted in buffer (20 mM HEPES, pH 7.4, 0.15 M NaCl, 1 mM DTT) to produce 12 concentrations, with the highest concentration adjusted based on the expected *K*_d_ for different protein variants. Labeled RNA was diluted to 10 nM in buffer (20 mM HEPES, pH 7.4, 0.15 M NaCl, 1 mM DTT, 0.1% (v/v) Tween20, 20 µg mL^-1^ yeast tRNA (Ambion^TM^). The protein and RNA solution was mixed at a 1:1 (v/v) ratio and incubated at room temperature for 30 min. To measure DND1 dose response in the presence of NANOS3, NANOS3 protein was added to the RNA solution to 4 µM before mixing with DND1. The samples were then loaded into MonolithNT.Automated capillary chips, and the measurements were performed at 25°C using a MonolithNT Automated instrument with picoRed excitation (NanoTemper Technologies). Medium MST power was applied. Data from three independently pipetted measurements were analyzed with MO.Affinity Analysis software (NanoTemper Technologies), and *K*_d_ values were derived using the signal from an MST- on time of 0.5 s. *K*_d_ values of NANOS3 were determined similarly by measuring the dose response of NANOS3 in the absence or in the presence of 4 µM DND1 protein. The reported errors are the confidence interval of the *K*_d_ from data fitting. The *K*_d_ is within the given range with a confidence of 68%.

### Crosslinking and mass spectrometry analysis

A protein solution with 7 µM DND1 1-230 and 11 µM NANOS3 ZF+C was prepared in a buffer of 20 mM HEPES, pH 7.4, 0.15 mM NaCl, 1 mM DTT. Two sets of samples were processed in parallel: one set with no RNA and one set with 10 µM *CDK1*-wt RNA. Protein or protein-RNA solution was divided into 50 µL aliquots. BS3 crosslinker solutions were prepared in water at 10, 5, 2.5, and 1 mM. 5 µL of BS3 crosslinker (ThermoFisher Scientific) was added to the samples at four different final concentrations (1, 0.5, 0.25, and 0.1 mM), and no BS3 was added to a fifth sample. Samples were incubated at room temperature for 30 min. Reactions were stopped by adding 3 µL 1 M Tris pH 8.0 to a final concentration of 50 mM and incubating at room temperature for 15 min. Samples were visualized by running 12 µL samples on an SDS-PAGE gel and staining with SimplyBlue SafeStain (Invitrogen) (Extended Data Fig. 4h). The crosslinked sample containing RNA with 0.1 mM BS3 was subject to mass spectrometry analysis.

Samples were digested with 100 ng trypsin (10 μL of Promega modified porcine trypsin at 10 ng μL^-1^ in ammonium bicarbonate pH 6.8) via direct addition of the trypsin to the samples. Digests were allowed to proceed overnight at room temperature. Peptide digests were transferred to 1.2 mL sample vials and placed in an autosampler at 4°C. Digests were analyzed by LC-MS/MS on an Orbitrap Ascend mass spectrometer (ThermoFisher Scientific) interfaced with a nanoVanquish UPLC system (ThermoFisher Scientific) equipped with a Easy-Spray™ PepMap™ Neo 2 μm C18 75 μm X 150 mm analytical column and a PepMap™ Neo 5 μm C18 300 μm X 5 mm cartridge column for trapping. 5 μL of peptide digest was injected onto the column. After a 5 min hold at 3% B, peptides were eluted by using a linear gradient from 3% to 45% over 105 min, followed by a step to 95% and a 10 min hold at 95%. The system flow rate was 350 nL min^-1^ and solvent A was 0.1% formic acid in water and solvent B was 0.1% formic acid in acetonitrile. Prior to analyses, the instrument was calibrated with Pierce™ FlexMix™ Calibration Solution (ThermoFisher Scientific). The mass spectrometer was operated in positive ion mode with a spray voltage of 2000 volts, and the Orbitrap was used for both MS1 (120,000 resolution, AGC target) and MS2 (30,000 resolution), and HCD for fragmentation.

Peak lists were generated from the LC/MS data using Mascot Distiller (Matrix Science, Boston, MA, U.S.A.), and the resulting peak lists were searched using the Batch-Tag Web function of the ProteinProspector web-based software developed by the UCSF Mass Spectrometry Facility. The MGF file was searched against sequences for the recombinant proteins and peptides by employing the User Protein Sequence field with other search parameters, including: tryptic specificity and 3 missed cleavages; precursor charge range of 2, 3, 4, and 5; monoisotopic values; parent mass tolerance of 10 ppm and fragment mass tolerance of 50 ppm; oxidation of methionine and incorrect monoisotopic assignment as a variable modifications; and in the Crosslinking field, the Link Search Type was defined as DSS. The putative cross-linked peptide output was triaged by limiting the mass error of putative cross-links to two standard deviations from the average error (9.09 ppm for this data set); requiring a Score Difference value >0 except for the cases of intermolecular cross-links of identical peptides or where one peptide was 6 amino acid residues or less in length; total expectation values below 1 x 10^-4^. Residue 0 of DND1 and residue 54 of NANOS3 are serine residues encoded by the BamHI restriction enzyme site used for subcloning.

### Animal Experiments

All animal experiments were conducted in accordance with the National Institutes of Health (NIH) Animal Care and Use regulations and were approved by the Animal Care and Use Committee (ACUC) of the National Institute of Arthritis and Musculoskeletal and Skin Diseases (NIAMS) under the ASP number A023-09-03. C57BL/6J mice were obtained from The Jackson Laboratory (JAX: 000664). The *Cdk1* N3-DRE mutant mouse line was generated in this study by Dr. Chengyu Liu at the Transgenic Core Facility of the National Heart, Lung, and Blood Institute (NHLBI) using CRISPR/Cas9-mediated genome editing (see *Generation of the Cdk1 N3-DRE mutant knock-in mouse line*).

### Generation of the Cdk1 N3-DRE mutant knock-in mouse line

The *Cdk1* N3-DRE-mutant knock-in mouse line was generated using the CRISPR/Cas9 method as previously described^74^. Briefly, a single guide RNA (sgRNA, GAATTATATTTAAATTCATA) was designed to cut in the 3′ UTR region of the mouse *Cdk1* gene, around the seven bp to be mutated, which was chemically synthesized by Synthego. A single-strand oligonucleotide donor was purchased from IDT (CTGTCATCTGGACTTTTCTTAATTTCCTACGTATAACTTAATTAACATGTAAATATTA TTCCAT**<u>GCC</u>**A**<u>C</u>**T**<u>C</u>**TAAATATAATTCTGTATATGTGCAGATGTCACTGTGGTGGCTGT TAATTACTATAACACAAGTG; the five underlined nucleotides in bold are mutated). The sgRNA was first incubated with Cas9 protein (IDT) to form Cas9-sgRNA RNP complex, which was then co-electroporated with the oligo template into zygotes collected from C57BL/6 mice using a Nepa21 electroporator (Nepa Gene Co.) following previously described procedures^75^. The electroporated zygotes were cultured overnight in KSOM medium (Millipore Sigma) at 37°C with 6% CO_2_. Those embryos that reached the 2-cell stage of development were implanted into the oviducts of pseudopregnant surrogate mothers (Swiss Webster mice from Charles River). Mice born to the foster mothers were genotyped by PCR amplification followed by Sanger sequencing to identify mice with the desired nucleotide changes. The PCR primers used for genotyping are listed in Supplementary Table 4. Heterozygous Cdk1 N3-DRE mutant mice (Cdk1^wt/mut^) were backcrossed to C57BL/6J mice. Mendelian ratios were assessed from litters obtained by intercrossing Cdk1^wt/mut^ mice between the second and fourth backcross generations. Statistical significance of deviations from expected Mendelian ratios (1:2:1) was evaluated using a Chi-square test. For immunofluorescence studies, Cdk1^wt/mut^ mice backcrossed for more than four generations were intercrossed to obtain homozygous mutants.

### Immunofluorescence of migrating PGCs in E8.5 embryos

For analysis of migrating primordial germ cells (PGCs), wild-type (WT) embryos were collected from C57BL/6J WT × WT crosses, and homozygous Cdk1^mut/mut^ embryos were obtained from Cdk1^mut/mut^ × Cdk1^mut/mut^ crosses. Embryos were harvested at E8.5, with the day of plug observation considered E0.5. Embryos were fixed in 4% paraformaldehyde (PFA) in PBS at 4°C for 1 h, washed three times for 5 min each with 1× PBS, and dehydrated through an ethanol series (30%, 50%, 70%, 100%; 5 min each). Paraffin embedding and sagittal sectioning were performed by HistoServe Inc. (https://www.histoservinc.com). Paraffin sections were deparaffinized in xylene (30 min), rehydrated through a descending ethanol series (100%, 70%, 50%, 30%; 5 min each), and washed in PBS (5 min). Antigen retrieval was performed using citrate buffer (Thermo Fisher Scientific, 005000) in a pressure cooker on high-pressure mode for 10 min, as described in^76^. Slides were washed twice for 5 min in PBST (PBS + 0.1% Tween-20), followed by blocking for 1 h at room temperature and overnight incubation at 4°C with primary antibodies using the M.O.M. Immunodetection Kit (Vector Laboratories, BMK-2202) according to the manufacturer’s instructions. Following three 5 min washes in PBST, slides were incubated with fluorescent secondary antibodies for 30 min at room temperature. After three additional PBST washes (5 min each), nuclei were counterstained with DAPI (100 ng mL^-1^ in PBST) for 7 min, followed by a final series of three PBST washes.

Coverslips were mounted using the Vectashield antifade mounting medium (Vector Laboratories, H-1000) and allowed to cure. Imaging was performed using a Leica TCS SP8 X confocal microscope employing either a HC PL Apo CS2 20X/0.75NA air lens or a HC PL Apo CS2 63X/1.4NA oil immersion lens (Leica) driven by the LAS X software (Leica). Large field embryo micrographs were obtained using the Leica tile-scanning modality (Navigator) in standard confocal resolution with image format set at 1,024x1,024 pixels XY, zoom 0.75, and pinhole set to 1 Airy Unit (AU). High magnification insets were captured using the Leica Lightning modality to achieve sub-diffraction resolution, with image format set at 2,048x2,048 pixels XY, zoom set to 1, and pinhole size 0.4-0.5 AU, using the ‘Adaptive’ deconvolution modality. Illumination of samples was achieved in frame mode to avoid crosstalk between channels with laser lines set to 405nm from a UV solid-state laser, and 488nm and 568nm using the Leica White Light Laser (WLL). Images were acquired as .lif or .lof and then exported as .tiff to be analyzed using Fiji (v2.16.0)^77^. Primary and secondary antibodies used are listed in Supplementary Table 4.

### Statistical analysis

Data are presented in figure legends as mean ± s.d. from three or four independent biological replicates. Statistical analyses were performed in R using two-sided Student’s t-tests for cell cycle and western blot quantification, and two-sided Wald tests for DESeq2 differential expression analysis. Further details are provided in the Source Data.

## Extended Data Figures and Legends

**Extended Data Figure 1.**
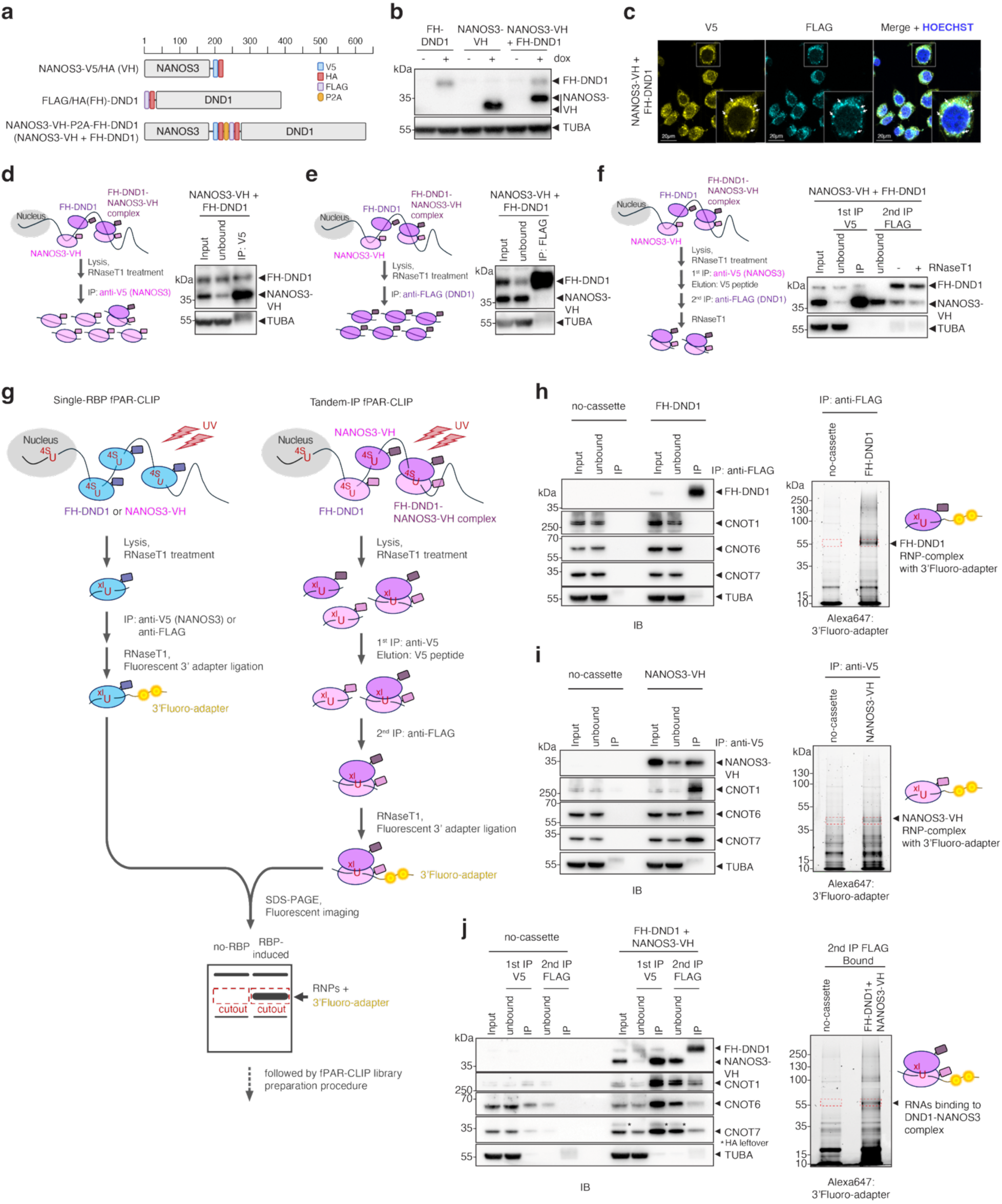
Tandem PAR-CLIP to identify sites co-occupied by DND1 and NANOS3. **(a)** Flp-In HEK293 cell lines used in this study express doxycycline-inducible gene cassettes for NANOS3-VH, FH-DND1, and NANOS3-VH-P2A-FH-DND1. **(b)** Immunoblot of cell lines from (a), with or without doxycycline treatment. Top blot probed with anti-HA antibody; bottom panel shows loading controls probed with anti-TUBA. (c) Representative immunofluorescence images showing cytoplasmic localization of co-expressed DND1 and NANOS3. Left image stained with anti-V5 antibody (NANOS3); middle panel, anti-FLAG antibody (DND1). Right image, merged, including DAPI stain to highlight the nucleus. **(d–f)** IP experiments with cell lines from (a). Antibodies and IP workflow shown in scheme to the left. Blots were probed with anti-HA antibody staining DND1 and NANOS3 (indicated). TUBA served as loading control. **(g)** Schematic of fPAR-CLIP to identify binding sites of individual RBPs (left) and Tandem PAR-CLIP (right) to identify binding sites co-occupied by DND1 and NANOS3. **(h- j)** Cross-linked, fluorescent adapter–ligated RNPs recovered by FH-DND1 (h) and NANOS3-VH (i) and Tandem PAR-CLIP experiments (j). Representative western blots (left) were probed with the indicated primary antibodies. Fluorescence images (right) show detection of RNA crosslinked to RBP and ligated to a 3′ adapter conjugated with Alexa Fluor 647.

**Extended Data Figure 2.**
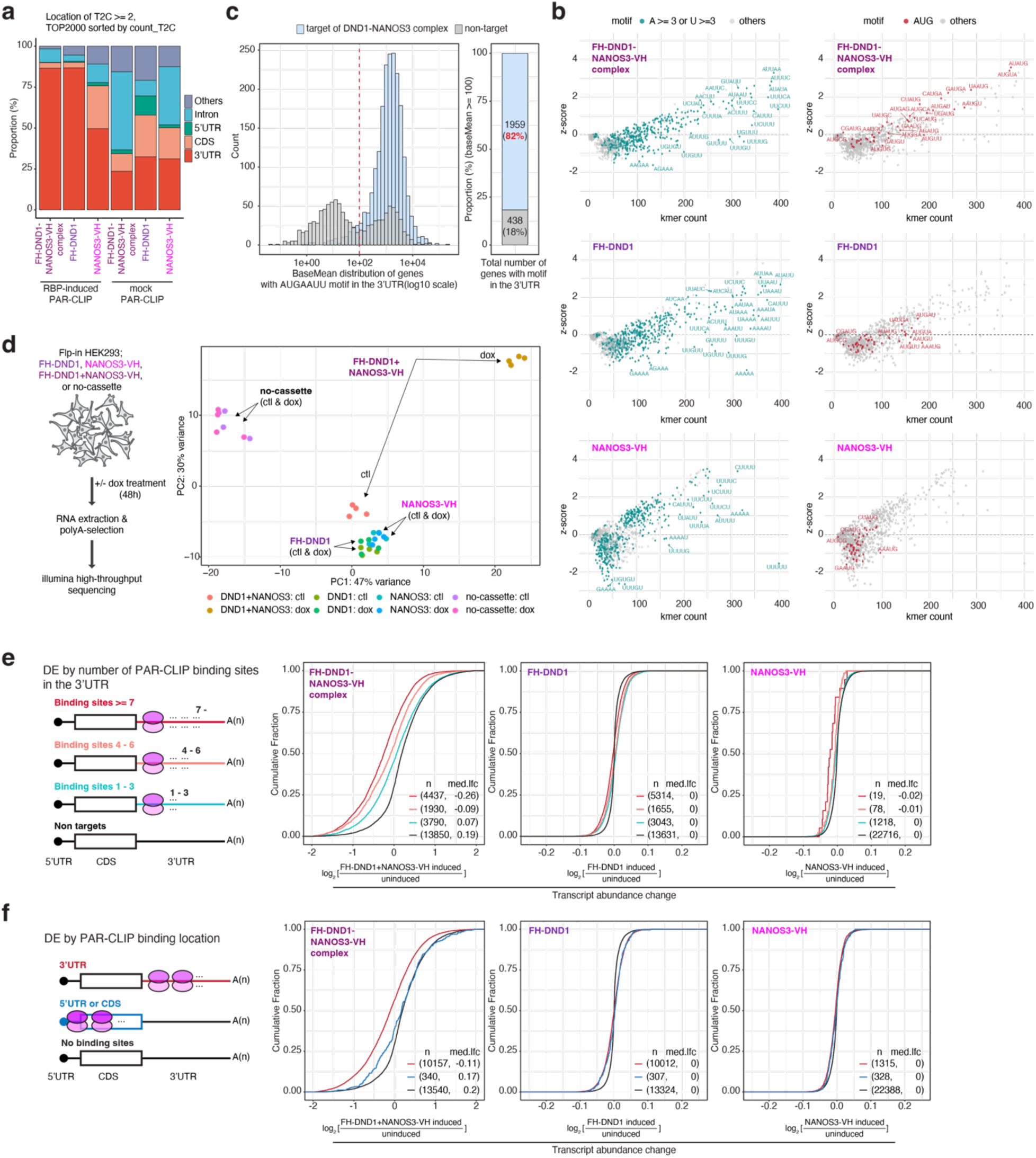
The DND1/NANOS3 complex binds the N3-DRE in mRNA 3′ UTRs and destabilizes them. **(a)** Stacked bar graph of PAR-CLIP and Tandem PAR-CLIP experiments, counting binding site location within transcripts. Biological replicates were merged after showing that they were highly similar (Fig. 1a). **(b)** k-mer analysis counting the number of occurrences for each possible 5-mer in our PAR-CLIP experiments for DND1 or NANOS3, and the Tandem PAR- CLIP mapping the DND1-NANOS3 complex sites. (left) 5-mers containing ≥3 A’s or ≥3 U’s are shown in cyan. (right) AUG-containing 5-mers are indicated in red. Y-axis documents the Z-score for each 5-mer. **(c)** (Left) Histogram of expression (DESeq2 baseMean) of transcripts containing the N3-DRE bound by the DND1-NANOS3 complex (light blue) or not bound (gray). The dashed red line indicates baseMean = 100. (Right) Proportional bar plot of genes with baseMean ≥ 100, showing DND1-NANOS3 targets (light blue) and non-targets (gray). (**d**) Quality control of RNA- seq samples. (left) Workflow of sample preparation from Flp-In HEK293 cell lines (FH-DND1, NANOS3-VH, FH-DND1+NANOS3-VH, and no-cassette). (right) PCA plot of all replicates. Each condition included four replicates, which clustered together. **(e)** Cumulative distribution function of transcript abundance changes upon induction of DND1-NANOS3 (left), DND1 alone (middle), or NANOS3 alone (right). Transcripts were binned according to number of binding sites on transcript. **(f**) Same as in (e), but transcripts were binned according to whether they were bound in the 3′ UTR, CDS, 5′ UTR, or not bound.

**Extended Data Figure 3.**
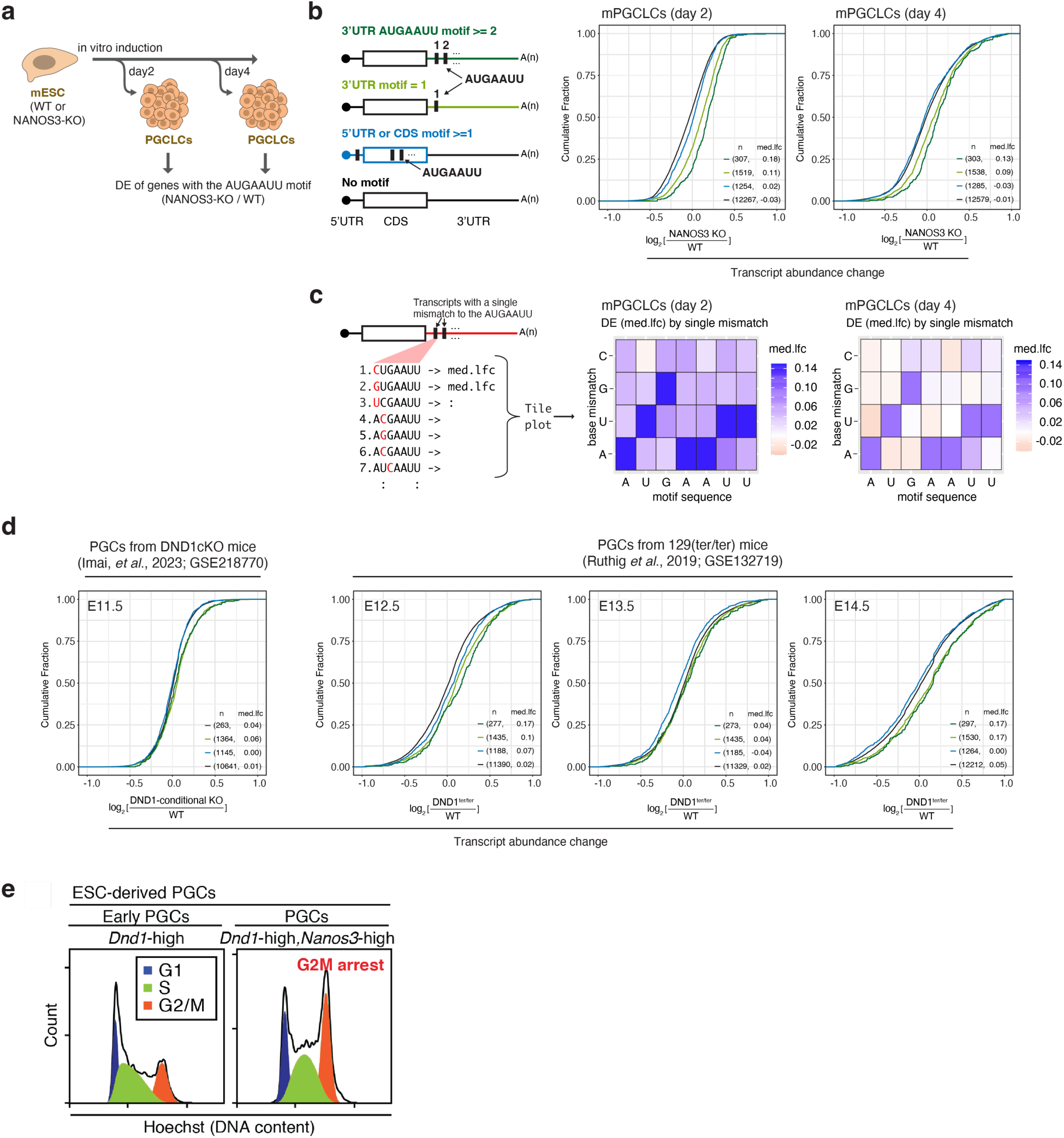
The DND1/NANOS3 complex suppresses mRNAs containing N3- DREs in their 3′ UTR in PGCs and PGCLCs. **(a)** Schematic of PGCLC induction from WT or Nanos3-KO mouse embryonic stem cells (mESCs). PGCLCs were collected for RNA sequencing at two or four days after induction. **(b)** Cumulative distribution function of transcript abundance changes in WT PGCLCs and NANOS3-KO PGCLCs. Transcripts were binned by presence or absence of the N3-DRE in their 3′ UTR (≥2 or 1), or 5′ UTR/CDS. **(c)** Tolerance of the N3-DRE to single-nucleotide variation. Panel definitions and schematic as in Fig. 1f. **(d)** Same as in (b), but for FACS-sorted PGCs from different developmental stages (indicated above). (**e**) Cell cycle analysis from developing PGCLCs, sorted according to whether they were expressing high levels of Dnd1 alone (left plot) or high levels of both Dnd1 and Nanos3.

**Extended Data Figure 4.**
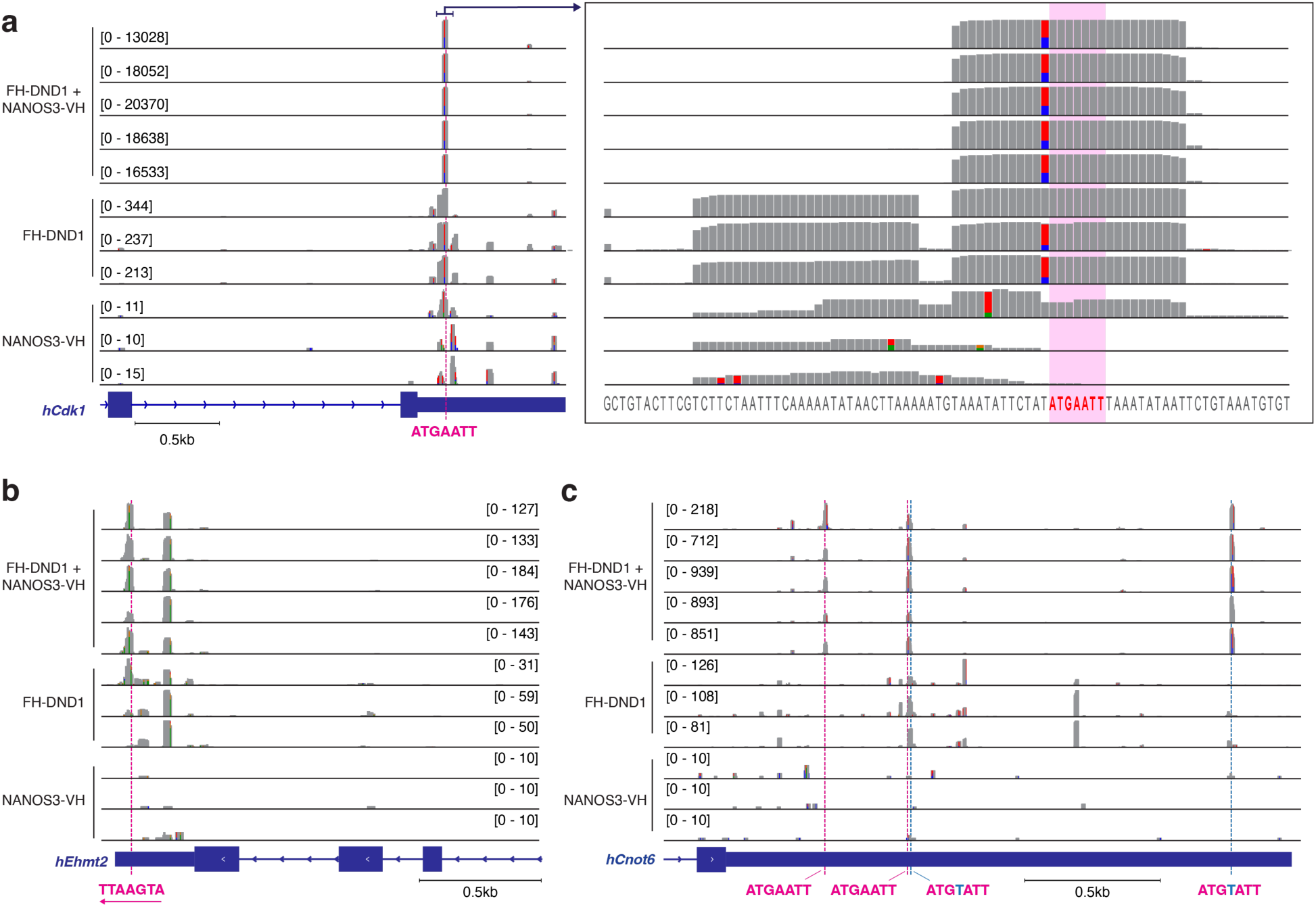
Binding sites of DND1, NANOS3, and the DND1-NANOS3 complex on human *CDK1*, *EHMT2* and *CNOT6* gene. (**a-c**) IGV track of PAR-CLIP and Tandem PAR-CLIP replicates around the 3′ UTR of *CDK1* (a), *CNOT6* (b), and *EHMT2* (c). Colored bars in each track indicate positions with a mutation ratio ≥20%. The colored dashed line marks the position of N3-DRE (pink), and its single mismatch (light blue). The right panel in (a) shows an enlargement of the indicated area, with base identity shown at the bottom and N3-DRE highlighted in red.

**Extended Data Figure 5.**
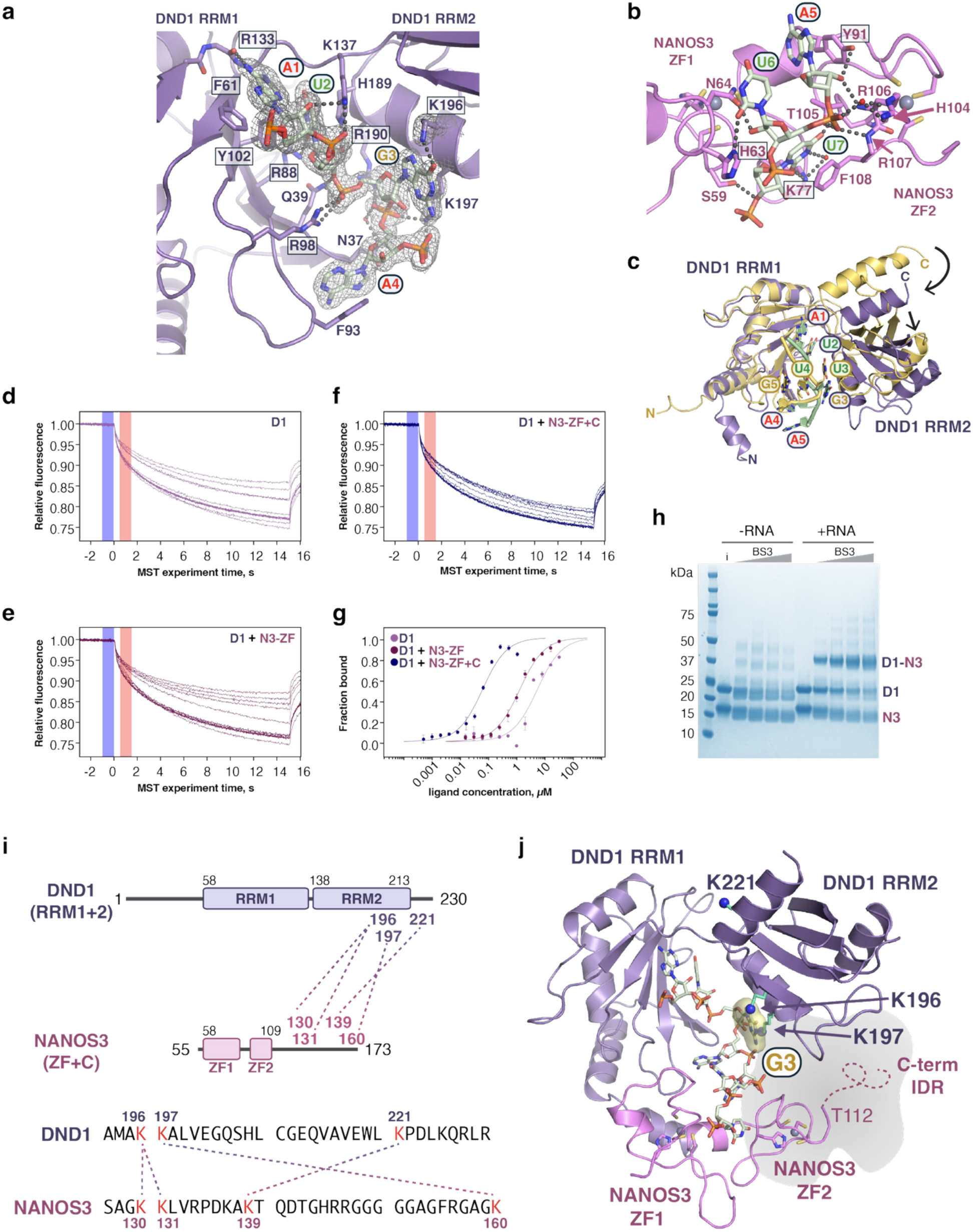
High-affinity DND1–NANOS3 RNA binding and G3 recognition require the NANOS3 C-terminal IDR. (**a**) DND1 interactions with the 5′ N3-DRE half-site A1- A4. A composite omit 2mFo-DFc electron density map contoured at 1σ is superimposed on the RNA and K196 and K197 side chains. (**b**) NANOS3 interactions with the 3′ N3-DRE half-site A5- U7. (**c**) Superposition of DND1 bound to N3-DRE (purple) and solution structure of DND1 bound to AU-rich RNA^42^ (yellow, PDB ID 7q4l). The individual RRM domains are structurally similar, with RMSD 1.5 Å for RRM1 and 1.7 Å for RRM2. However, alignment of RRM1 in this figure illustrates the different relative positions of RRM2. The RNA sequence in the NMR structure does not match the N3-DRE, however, nts A1 and U2 are common to both structures. The remaining RNA structures are divergent. (**d-f**) MST traces from one measurement of (d) DND1, (e) DND1 + NANOS3 ZF, or (f) DND1 + NANOS3 ZF+C binding to CDK1 N3-DRE RNA. Blue bars indicate F_cold_ at 0 s before the IR-laser was turned on, and red bars indicate F_hot_ at 0.5 s after the laser was turned on. Normalized fluorescence F_norm_ is defined as F_hot_/F_cold_. (**g**) MST analysis of binding to 5′-Cy5-labeled CDK1 N3-DRE RNA. Data were fit using the *K*_d_ model included in the MO.Affinity Analysis software (NanoTemper). The mean fraction of bound RNA calculated from the change in F_norm_ from three independently pipetted replicates is plotted vs the concentration of protein. (**h**) SDS-PAGE gel of BS3-crosslinked samples of DND1–NANOS3 ZF+C in the absence (left) or presence (right) of N3-DRE RNA. Uncrosslinked input sample (i) indicates the migration of DND1 1-230 (26 kDa) and NANOS3 ZF+C (13 kDa). In the +RNA lanes, a band corresponding to the expected size of DND1–NANOS3 ZF+C (39 kDa) is detected (D1–N3), whereas no complex is detected in the -RNA lanes. (**i**) Mass spectrometry analysis of BS3-crosslinked DND1–NANOS3 ZF+C + RNA identifies crosslinks between the NANOS3 C-terminal IDR and the DND1 C-terminal region that bind N3-DRE RNA G3. Crosslinked lysine residues are indicated by dotted lines. Intramolecular crosslinks and crosslinks between N-terminal amine groups are not shown. A full list of crosslinked peptides is provided in Supplementary Table 3. (**j**) Locations of crosslinked DND1 lysine residues indicated on ribbon diagram of DND1–NANOS3 ZF bound to CDK1 N3-DRE RNA. DND1 lysine residues are shown as sticks with blue nitrogen spheres. The G3 base is shown with a yellow surface, and the NANOS3 C-terminal IDR, not visible in the crystal structure, is represented by a dashed line and grey cloud to indicate its proximity to the DND1 RRM2. T112 is the last ordered residue in NANOS3.

**Extended Data Figure 6.**
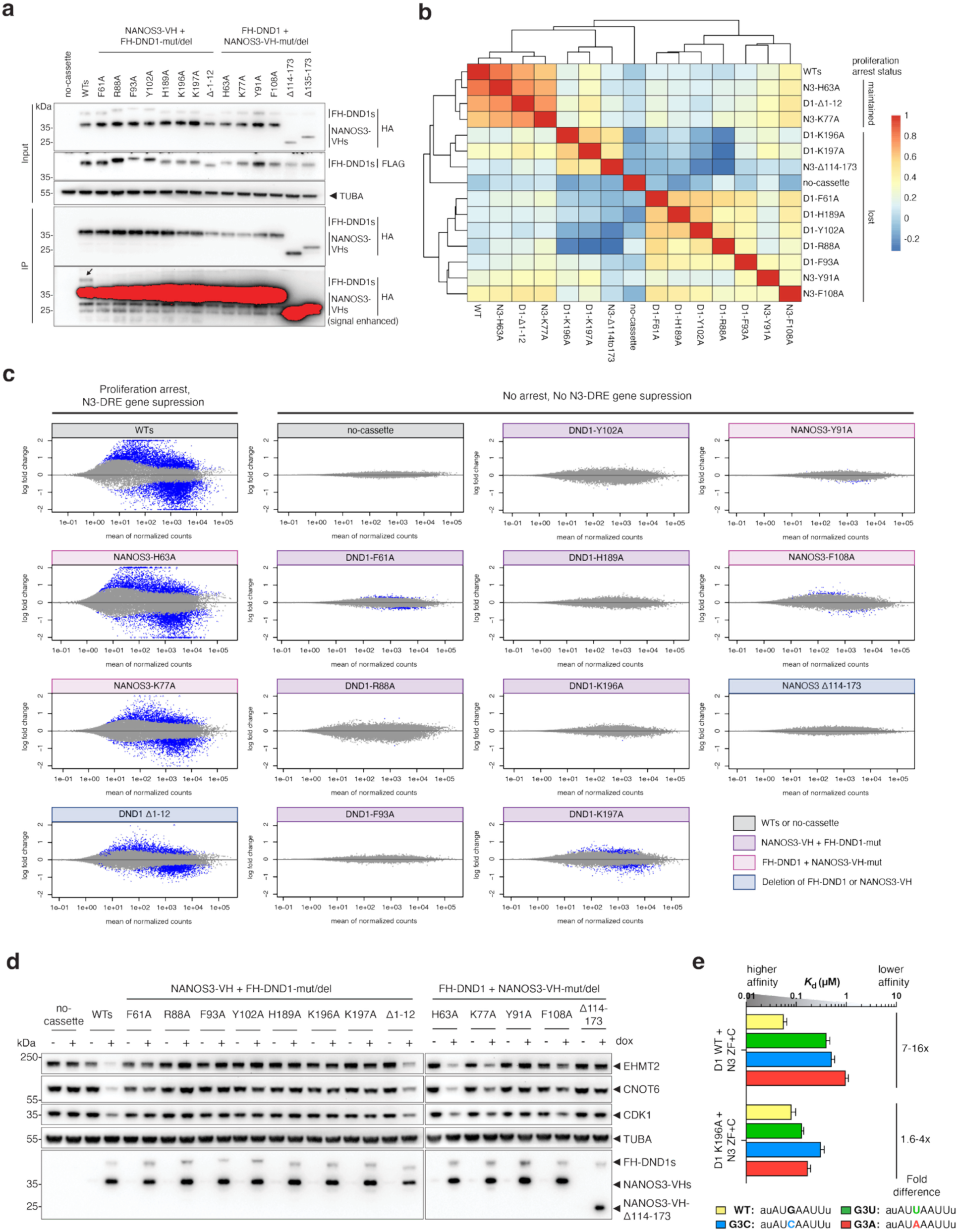
RNA-binding mutants of DND1 and NANOS3 are not functional. **(a)** Co-IP of DND1 and NANOS3 mutant or deletion constructs. The mutations (see also Fig. 4b) are indicated. Co-IPs were performed with anti-V5 antibody (NANOS3) in two independent experiments. The black arrow marks co-immunoprecipitated DND1. Blots were probed with the indicated antibodies. **(b–c)** Classification of mutant or deletion cell lines based on effects on differential expression. (b) Correlation heatmap of DESeq2 log₂ fold changes across DND1 and NANOS3 mutant constructs, WTs (DND1-WT + NANOS3-WT), and no-cassette cells. Mutants that maintain proliferation arrest in Fig. 4d-f cluster with WTs, indicating preserved function. (c) MA plot showing similar trends, with additional non-phenotypic mutations (e.g. DND1-F61A, - K197A, NANOS3-F108A) displaying weaker effects. Blue points denote significant genes (adjusted p-values < 0.05). **(d)** Representative Western blot showing loss of suppression of N3- DRE targets (CDK1, CNOT6, EHMT2) by most DND1-NANOS3 mutants or deletion. **(e)** DND1 K196A loses selectivity for the N3-DRE G3 nucleotide *in vitro* in MST binding assays. *K*_d_ values were determined from 3 independently pipetted measurements.

**Extended Data Figure 7.**
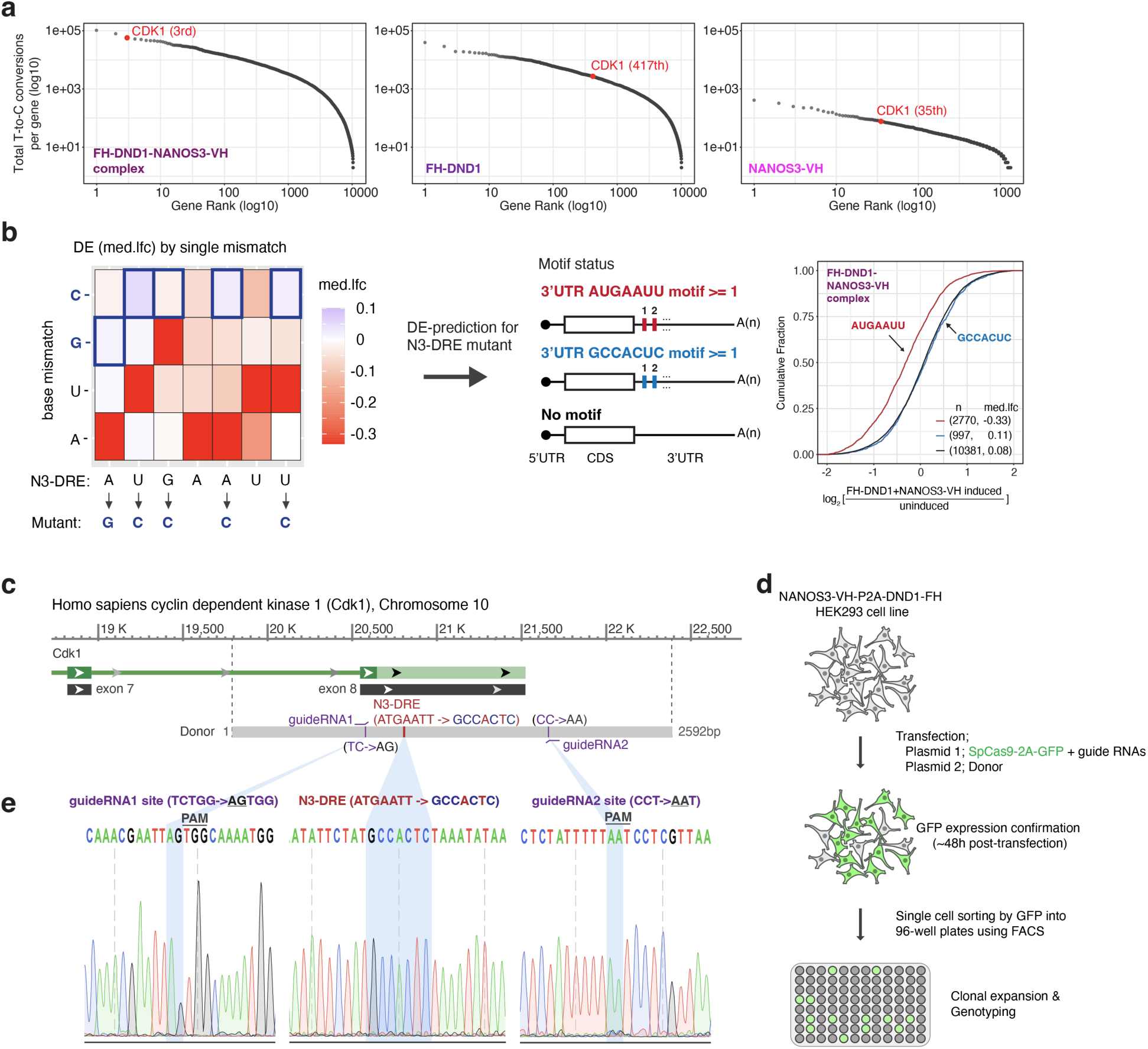
Design and generation of CDK1 N3-DRE mutant cells. **(a)** Ranking of PAR-CLIP targets by the total number of T-to-C conversions per gene, shown for targets of DND1-NANOS3 complex (left), DND1 alone (middle), and NANOS3 alone (right). **(b)** Design of N3-DRE mutations based on DE prediction. Tile plot analysis (Fig. 1f) was used to identify positions and nucleotides capable of abolishing suppression. The GCCACUC substitution was selected as the N3-DRE mutation (left), and CDF analysis (right) confirmed that genes carrying this mutated motif were no longer suppressed by DND1 and NANOS3 co-expression. **(c)** Double-stranded DNA donor designed to introduce mutations within the N3-DRE in the 3′ UTR of CDK1. Additional mutations were incorporated into the guide RNA binding sites to prevent recurrent Cas9 cutting. **(d)** Workflow for generating CDK1 N3-DRE mutant cells in Flp-in NANOS3-VH- P2A-FH-DND1 HEK293 cells. **(e)** Genotyping by Sanger sequencing at the mutation site. (Left) guide RNA1 site; (middle) N3-DRE; (right) guide RNA2 site.

**Extended Data Figure 8.**
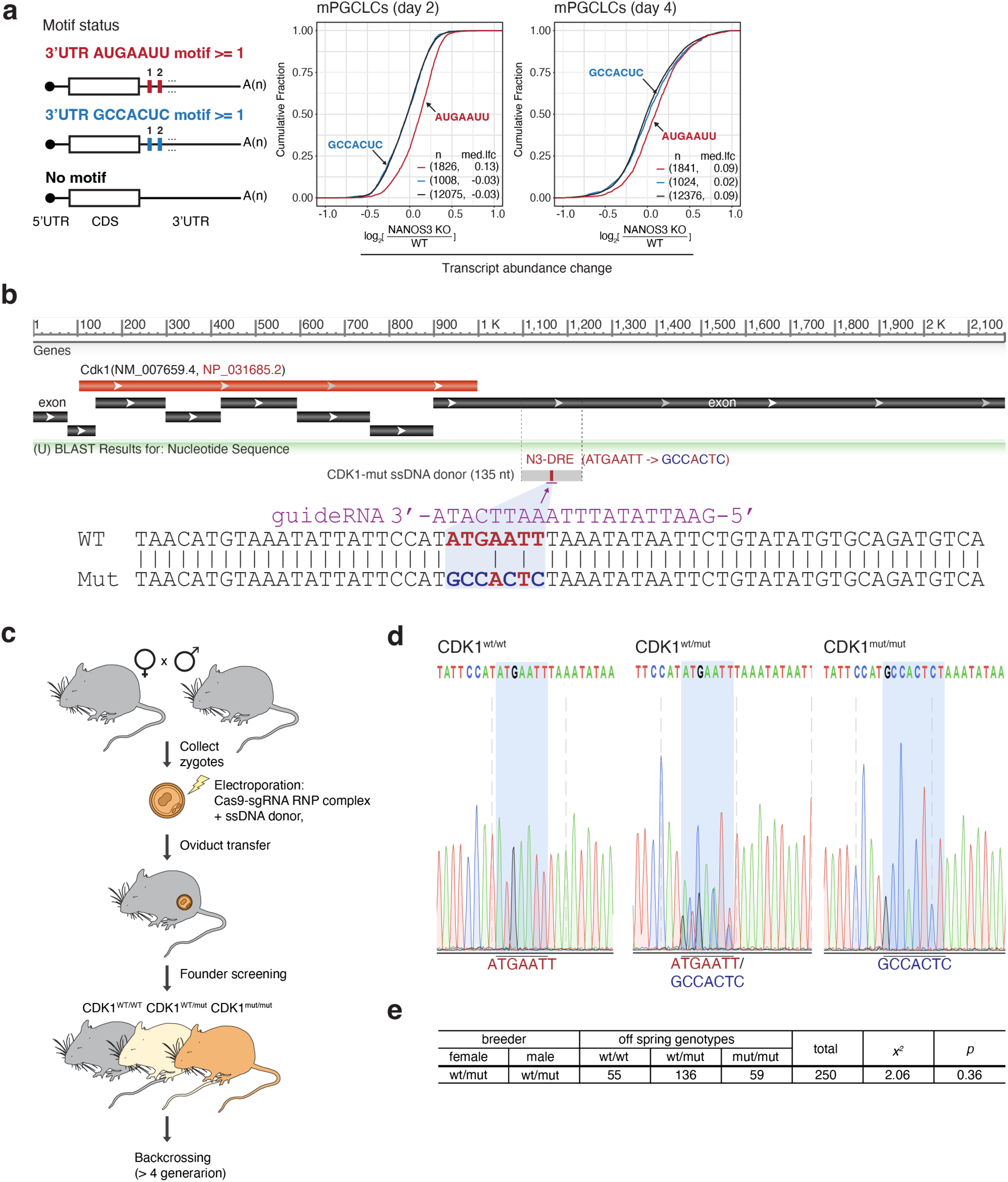
Generation of the Cdk1 N3-DRE mutant knock-in mouse line. (**a**) DE analysis of genes bearing the N3-DRE or its mutant (GCCACUC) in WT and *Nanos3*-KO PGCLCs at days 2 and 4 post-induction. CDF plots indicate that transcripts containing the mutant motif are unaffected by *Nanos3* KO at either time point. **(b)** Single-stranded DNA donor designed to introduce mutations within the N3-DRE in the 3′ UTR of *CDK1*. Bottom: Highlighted view of the mutated positions in the donor sequence. **(c)** Workflow for generating *Cdk1* N3-DRE mutant mice. Cas9-sgRNA RNP complexes and donor DNA were electroporated into zygotes collected from C57BL/6 mice, followed by oviduct transfer. Founder mice were genotyped using PCR amplification followed by Sanger sequencing, and heterozygous *Cdk1* N3-DRE mutant mice (*Cdk1*^wt/mut^) were backcrossed to C57BL/6J. **(d)** Sanger sequencing of the mutation site. Left: WT, middle: heterozygous; right: homozygous. **(e)** Mendelian ratio from a heterozygous × heterozygous cross.

