## Supplementary_Fig. 1 for "The DND1–NANOS3 complex shapes the primordial germ cell transcriptome via a heptanucleotide sequence in mRNA 3′ UTRs"

### Supplementary Figure 1

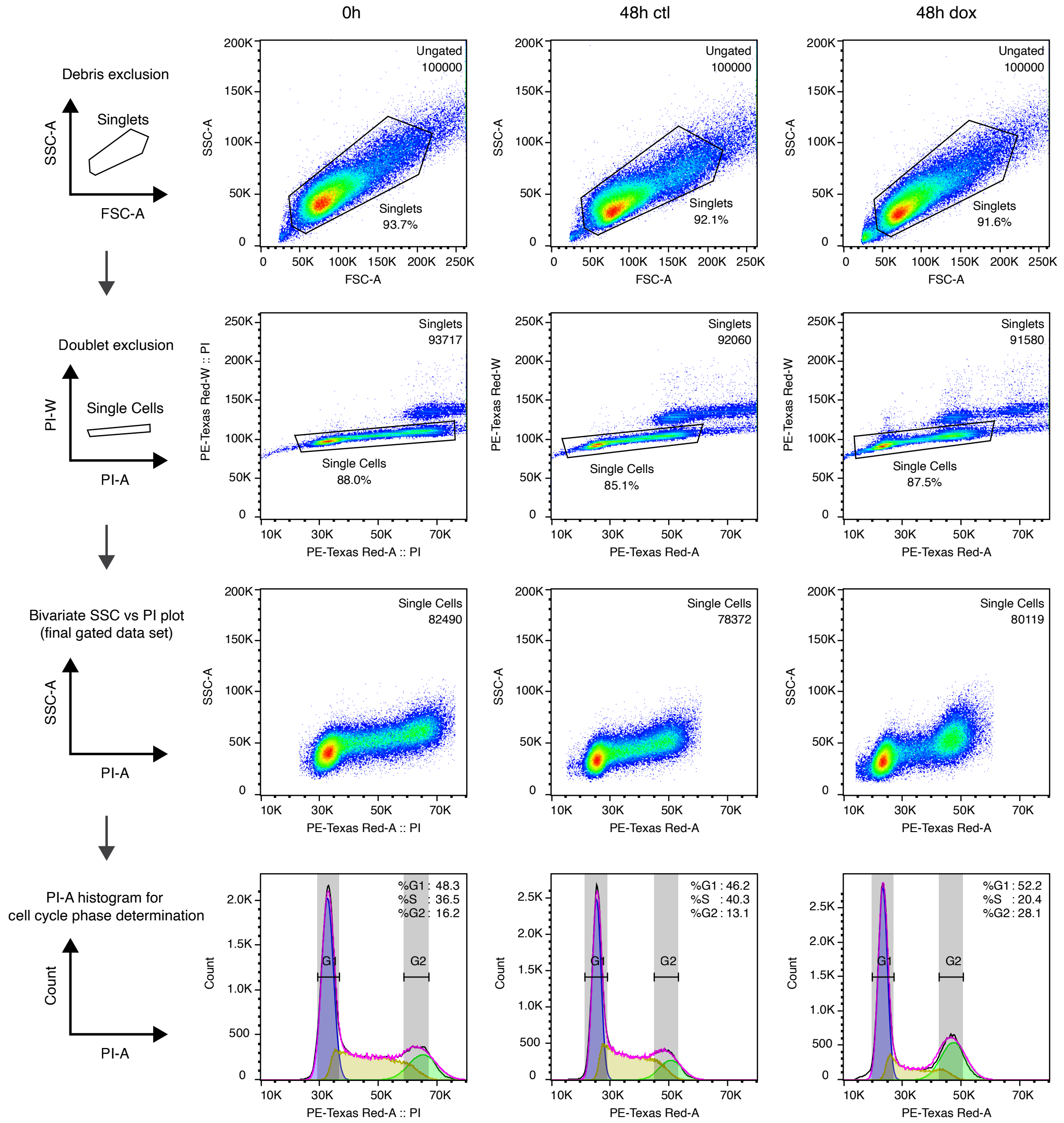


**Supplementary Figure 1: Gating strategy for cell cycle analysis relate to Fig. 2b.**
Representative data are shown for 0 h, 48 h control, and 48 h doxycycline-treated cells.
Fixed and stained cells were first gated on FSC-A versus SSC-A to exclude debris and on PI-A versus PI-W to exclude doublets. SSC-A versus PI-A plots with their corresponding PI-A histograms are shown as final gated dataset. DNA content histograms were used to determine the distribution of cells in G0/G1, S, and G2/M phases.
