## Supplementary Table 2 for "The DND1–NANOS3 complex shapes the primordial germ cell transcriptome via a heptanucleotide sequence in mRNA 3′ UTRs"

Supplementary Table 2 | RNA-binding affinities of DND1 and NANOS3

|  | **RBP1** | **+/- RBP2** | **RNA** | **RNA sequence** | ***K*_d_, μM**^1^ |
| --- | --- | --- | --- | --- | --- |
| 1 | DND1 1-230 | **-** | CDK1 | auAUGAAUUu | 5.1 ± 2.4 |
| 2 | DND1 1-230 | NANOS3 ZF | CDK1 |  | 1.2 ± 0.1 |
| 3 | DND1 1-230 | NANOS3 ZF+C | CDK1 |  | 0.06 ± 0.02 |
| 4 | DND1 1-230 | - | D1mut | auGCCAAUUu | 15 ± 4.7 |
| 5 | DND1 1-230 | NANOS3 ZF+C | D1mut |  | 15 ± 7.7 |
| 6 | DND1 1-230 | - | N3mut | auAUGACUCu | 6.4 ± 3.8 |
| 7 | DND1 1-230 | NANOS3 ZF+C | N3mut |  | 2.7 ± 1.1 |
| 8 | DND1 1-230 | - | D1-N3mut | auGCCACUCu | 11 ± 2.1 |
| 9 | DND1 1-230 | NANOS3 ZF+C | D1-N3mut |  | 12 ± 3.0 |
| 10 | DND1 1-230 | - | spacer | auAUGAcccAUUu | 8.2 ± 3.6 |
| 11 | DND1 1-230 | NANOS3 ZF+C | spacer |  | 14 ± 7.0 |
| 12 | DND1 1-230 | **-** | CDK1 G3 | auAUGAAUUu | 5.1 ± 2.4 |
| 13 | DND1 1-230 | - | G3U | auAUUAAUUu | 7.9 ± 3.0 |
| 14 | DND1 1-230 | - | G3C | auAUCAAUUu | 6.0 ± 2.7 |
| 15 | DND1 1-230 | - | G3A | auAUAAAUUu | 8.1 ± 3.5 |
| 16 | DND1 1-230 | NANOS3 ZF+C | CDK1 G3 | auAUGAAUUu | 0.06 ± 0.02 |
| 17 | DND1 1-230 | NANOS3 ZF+C | G3U | auAUUAAUUu | 0.40 ± 0.12 |
| 18 | DND1 1-230 | NANOS3 ZF+C | G3C | auAUCAAUUu | 0.51 ± 0.13 |
| 19 | DND1 1-230 | NANOS3 ZF+C | G3A | auAUAAAUUu | 0.97 ± 0.22 |
| 20 | DND1 1-230 | NANOS3 ZF | CDK1 G3 | auAUGAAUUu | 1.2 ± 0.1 |
| 21 | DND1 1-230 | NANOS3 ZF | G3U | auAUUAAUUu | 2.6 ± 0.8 |
| 22 | DND1 1-230 | NANOS3 ZF | G3C | auAUCAAUUu | 1.4 ± 0.5 |
| 23 | DND1 1-230 | NANOS3 ZF | G3A | auAUAAAUUu | 2.5 ± 1.0 |
| 24 | DND1 K196A | NANOS3 ZF+C | CDK1 G3 | auAUGAAUUu | 0.08 ± 0.03 |
| 25 | DND1 K196A | NANOS3 ZF+C | G3U | auAUUAAUUu | 0.13 ± 0.02 |
| 26 | DND1 K196A | NANOS3 ZF+C | G3C | auAUCAAUUu | 0.31 ± 0.09 |
| 27 | DND1 K196A | NANOS3 ZF+C | G3A | auAUAAAUUu | 0.17 ± 0.04 |
| 28 | DND1 F93A | NANOS3 ZF+C | CDK1 | auAUGAAUUu | 1.1 ± 0.4 |
| 29 | DND1 H189A | NANOS3 ZF+C | CDK1 |  | 0.16 ± 0.05 |
| 30 | DND1 13-230 | - | CDK1 | auAUGAAUUu | 3.7 ± 1.4 |
| 31 | DND1 13-230 | NANOS3 ZF | CDK1 |  | 4.0 ± 2.4 |
| 32 | DND1 13-230 | NANOS3 ZF+C | CDK1 |  | 0.11 ± 0.03 |
| 33 | NANOS3 ZF | **-** | CDK1 | auAUGAAUUu | Not detected |
| 34 | NANOS3 ZF | DND1 1-230 | CDK1 |  | 0.44 ± 0.08 |
| 35 | NANOS3 ZF+C | - | CDK1 | auAUGAAUUu | 109 ± 35 |
| 36 | NANOS3 ZF+C | DND1 1-230 | CDK1 |  | 0.12 ± 0.04 |

^1^ *K*_d_ values were determined from 3 independently pipetted measurements, and the reported errors are the confidence interval of the *K*_d_ from data fitting. The *K*_d_ is within the given range with a confidence of 68%.
